# Lysyl-phosphatidylglycerol promotes cell-to-cell interaction and biofilm formation of *Staphylococcus aureus* as a biofilm matrix component

**DOI:** 10.1101/2025.02.19.638968

**Authors:** Shinya Sugimoto, Keiichiro Hara, Yoshitaka Taketomi, Yuki Nagasaki, Chikara Sato, Makoto Murakami, Yuki Kinjo

## Abstract

Biofilms formed by *Staphylococcus aureus* contribute significantly to persistent infections and antibiotic resistance, driven by the unique composition of their extracellular matrix. While previous studies highlighted extracellular DNA, proteins, and polysaccharides as key components, the role of phospholipids in biofilm architecture remains underexplored. This study identifies extracellular phospholipids, including lysyl-phosphatidylglycerol (Lys-PG), phosphatidylglycerol, and cardiolipin (CL), as critical structural elements in *S. aureus* biofilms. Bacterial phospholipase A_1_ (PLA_1_) that cleaves the acyl ester bond at the *sn*-1 position of phospholipids effectively dispersed pre-formed biofilms and prevented new biofilm formation by hydrolyzing extracellular phospholipids, without affecting bacterial growth or exhibiting cytotoxicity. Microscopy analyses confirmed that PLA_1_ disrupts membranous nanostructures, including extracellular vesicles and nanofilaments, integral to biofilm stability. Lipidomic analysis revealed an enrichment of Lys-PG and CL in the biofilm matrix. Lys-PG promotes bacterial aggregation by acting as a molecular glue, mediated through electrostatic and hydrophobic interactions. Deletion of the *mprF* gene, responsible for Lys-PG synthesis, significantly impaired biofilm formation, confirming its essential role. These findings reveal a "moonlighting" function of phospholipids in biofilm architecture, providing insights into biofilm biology and presenting PLA_1_ as a promising tool for biofilm control.

**Significance:** Biofilms, dense bacterial communities, pose significant challenges across medical, industrial, and daily life contexts. Understanding their formation mechanisms is crucial for developing effective strategies against them. We investigated biofilm matrix components in *S. aureus* biofilms, focusing on phospholipids. Our study reveals the presence and significance of Lys-PG in the biofilm matrix, acting as a crucial factor in biofilm formation and maintenance. Through biochemical, lipidomic, and genetic analyses, we demonstrate the role of Lys-PG in facilitating cell-to-cell contacts, contributing to the robustness and thickness of *S. aureus* biofilms. These findings shed light on the physiological function of extracellular phospholipids in bacterial biofilms and suggest targeting Lys-PG and its synthetic mechanism as a promising strategy for biofilm control.

## INTRODUCTION

Biofilms are intricate communities of microorganisms that adhere to both biotic and abiotic surfaces, embedded within a matrix of self-produced extracellular polymeric substances such as proteins, polysaccharides, and nucleic acids (*1*). While biofilms offer benefits in environmental and industrial processes like carbon, nitrogen, and mineral cycling, wastewater treatment using active sludge, and food and beverage fermentation (*2*), they also pose significant challenges. These include contributing to metal corrosion in water systems, contaminating food supplies, and creating persistent biofilm-associated infections in healthcare (*3*). Biofilm formation on medical devices is especially concerning, as it can lead to chronic infections that are difficult to treat due to the tolerance of biofilms to antibiotics and evasion of the host immune system (*4*). Consequently, there is a growing need for innovative strategies to control and eliminate biofilms across various scientific and technological domains.

*Staphylococcus aureus* is a prominent opportunistic pathogen frequently linked to biofilm-associated infections, particularly in clinical settings (*5*). Around 30% of healthy individuals carry *S. aureus*, making them more susceptible to surgical site infections (*6*). Moreover, *S. aureus* is a well-known food-borne pathogen, responsible for infections transmitted through contaminated food (*7*). Understanding how *S. aureus* forms biofilms is crucial for developing effective approaches to address these biofilm-related challenges. In our previous research, we developed methods for isolating and characterizing matrix components from *S. aureus* biofilms using high-salt solutions (*8, 9*). Our studies revealed that extracellular DNA (eDNA) is a predominant component of the matrix in *S. aureus* biofilms, while only a few strains form biofilms through polysaccharide intercellular adhesin, also known as poly-N-acetylglucosamine (PNAG) (*10*). We have further demonstrated that various biofilm matrix components, including secreted proteins, cell wall-anchored proteins, PNAG, and environmental RNA (envRNA), work both cooperatively and redundantly (*11, 12*). However, the full composition of the biofilm matrix in *S. aureus* remains incompletely understood, and identifying previously unknown components could provide critical insights for new biofilm control strategies.

Phospholipids are well-known for their essential biological roles, including serving as key components of cell membranes, energy sources, and structural barriers. They also play crucial roles in signal transduction, metabolism regulation, and gene expression (*13*) (*14*). In *S. aureus*, the membrane phospholipid profile typically consists of 40–70% phosphatidylglycerol (PG) and 30–55% lysyl-phosphatidylglycerol (Lys-PG), both containing saturated and iso-branched fatty acids (*15*). Cardiolipin (CL), a diphosphatidylglycerol, is present in smaller amounts (4–9%) (*16*). Beyond their cellular roles, phospholipids are also components of extracellular nanostructures like membrane vesicles (MVs) and interbacterial nanotubes, which are involved in intercellular communication and host interactions (*17–19*). MVs, in particular, have been implicated in biofilm formation, as their hydrophobicity aids in forming intercellular junctions within biofilms (*20*). Despite these insights, the composition and function of extracellular phospholipids in the biofilm matrix of *S. aureus* remain largely unexplored.

In this study, we identified phospholipids in the matrix of *S. aureus* biofilms by lipidomic analysis. Treatment with bacterial phospholipase A_1_ (PLA_1_) prevented biofilm formation and dispersed pre-formed biofilms in various laboratory and clinical strains of *S. aureus*. Notably, Lys-PG, one of the abundant phospholipids identified in the biofilm matrix, played a significant role in promoting cell aggregation by acting as a molecular glue. These findings suggest that phospholipids have a previously unrecognized "moonlighting" function in biofilms, where they not only serve their traditional roles in cell membranes but also play a critical structural role in biofilm architecture.

## RESULTS

### Membranous Extracellular Nanostructures in *S. aureus* Biofilms

Membranous extracellular nanostructures, such as nanotubes and MVs, have been observed in biofilms of various bacterial species (*17, 18, 21, 22*). These structures are believed to play important roles in biofilm formation and development. In this study, we aimed to determine whether similar extracellular nanostructures are present in *S. aureus* biofilms, using multiple microscopy techniques.

First, we employed confocal laser scanning microscopy (CLSM) in combination with the iCBiofilm method, a recently developed biofilm-clearing technique to observe thick biofilms (*23*). We examined biofilms formed by the *S. aureus* strain MR23, which is known for producing robust and thick biofilms and has been frequently used in our previous studies (*8–12, 24*). The MR23 biofilm was stained with the lipophilic fluorescent probe FM1-43, which labels both the cytoplasmic membrane and extracellular membranous structures. As expected, MR23 formed a dense and rugged biofilm (Fig. 1A), consistent with prior reports (*11, 23*). Notably, at the base of the biofilm, numerous extracellular spherical nanostructures were observed (Fig. 1B).

**Fig. 1.**
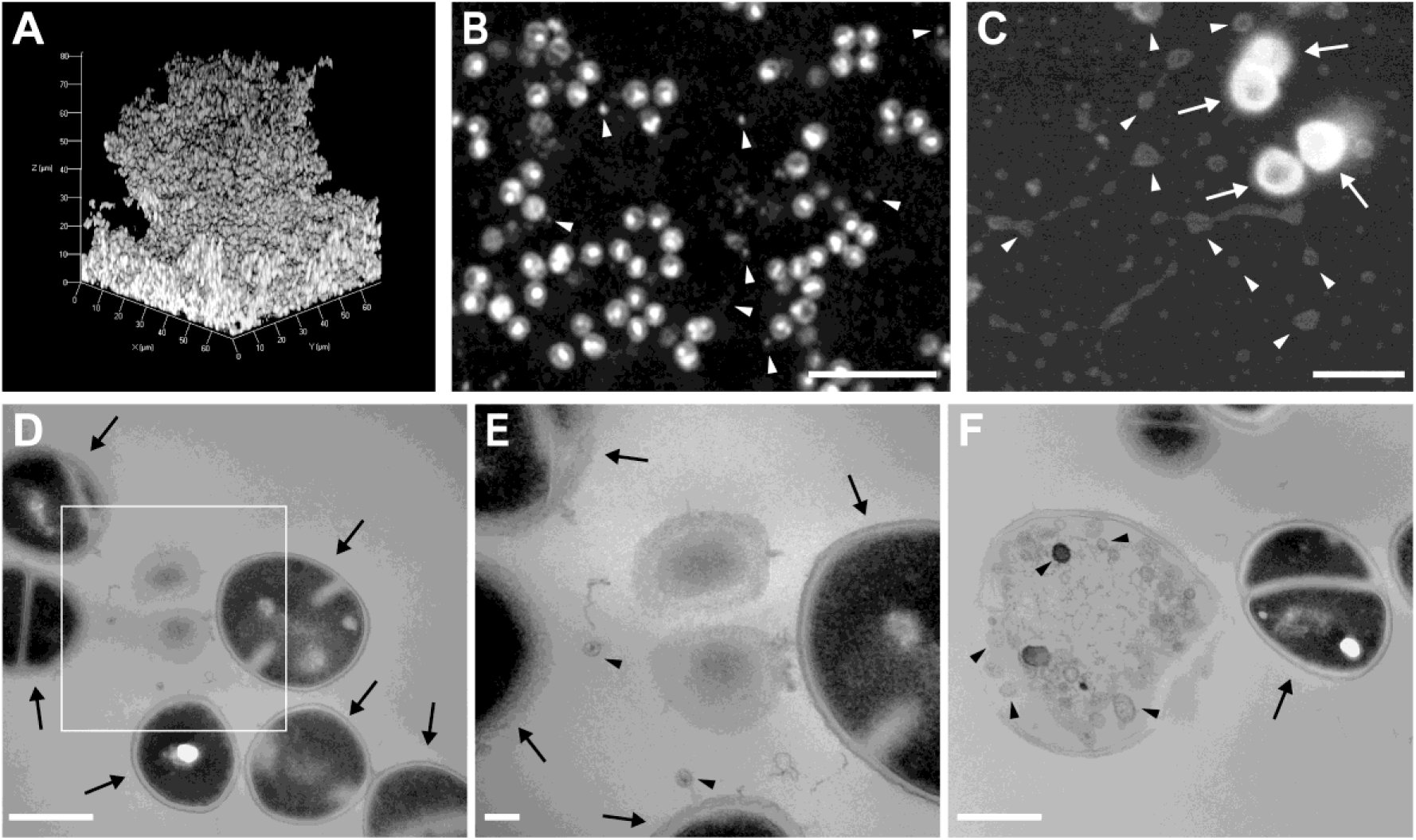
Detection of Phospholipid-Based Nanostructures in *S. aureus* Biofilms. (**A**) Three-dimensional CLSM visualization of the biofilm formed by *S. aureus* strain MR23. The biofilm was stained using the membrane-binding probe FM1-43 and optically cleared via the iCBiofilm method (*23*) prior to imaging. (**B**) High-magnification CLSM image showing the interface between the glass surface and the MR23 biofilm. (**C**) ASEM image of the biofilm-surface interface reveals spherical and filamentous nanostructures. (**D–F**) Thin-section TEM images of MR23 biofilm cells. (**E**) A higher magnification image of the white rectangle in **D**. (**D, E**) Membrane vesicles (MVs) are visible outside bacterial cells. (**F**) Dead cells display intracellular MVs. Arrows point to bacterial cells, while arrowheads indicate spherical nanostructures. Scale bars: 5 μm (**B**), 1 μm (**C**), 500 nm (**D**, **F**), and 100 nm (**E**).

Next, we used atmospheric scanning electron microscopy (ASEM) to visualize these extracellular nanostructures at higher resolution. ASEM allows for imaging biofilms in an aqueous environment, avoiding dehydration artifacts common with traditional electron microscopy techniques (*21, 25*). In agreement with the CLSM findings, we observed many extracellular spherical nanostructures of varying sizes (Fig. 1C). Additionally, filamentous structures connecting these spherical particles were also visualized by ASEM (Fig. 1C).

Finally, we employed transmission electron microscopy (TEM) for the highest resolution imaging of these membranous nanostructures. Thin sections of MR23 cells, harvested from the biofilm revealed the presence of extracellular MVs (Fig. 1, D and E), consistent with previous reports (*21*). Intracellular MVs were also observed in dead cells (Fig. 1F).

Together, these results indicate the presence of various membranous nanostructures, including phospholipid-containing vesicles and filaments, in *S. aureus* biofilms and that they could play some roles in biofilm stability and development.

### Phospholipases Inhibit Biofilm Formation and Disperse Mature Biofilms in Various *S. aureus* Strains

To investigate the role of extracellular phospholipids in biofilm development, mature biofilms formed by selected strains of *S. aureus*, which display distinct biofilm characteristics (*8–10*), were treated with various phospholipases. The cleavage sites for the phospholipases used in this study are shown in Fig. 2A. Phospholipases are lipolytic enzymes that hydrolyze phospholipids into free fatty acids and lysophospholipids. These enzymes are classified into four groups—A, B, C, and D—based on their catalytic reactions (*26*). Phospholipase A (PLA) enzymes cleave the acyl ester bond at either the *sn*-1 (phospholipase A_1_; PLA_1_) or *sn*-2 (phospholipase A_2_; PLA_2_) position of the glycerol backbone. Phospholipase C (PLC) cleaves before the phosphate group, releasing diacylglycerol and a phosphate-containing head group, while phospholipase D (PLD) cleaves after the phosphate, yielding phosphatidic acid and an alcohol.

**Fig. 2.**
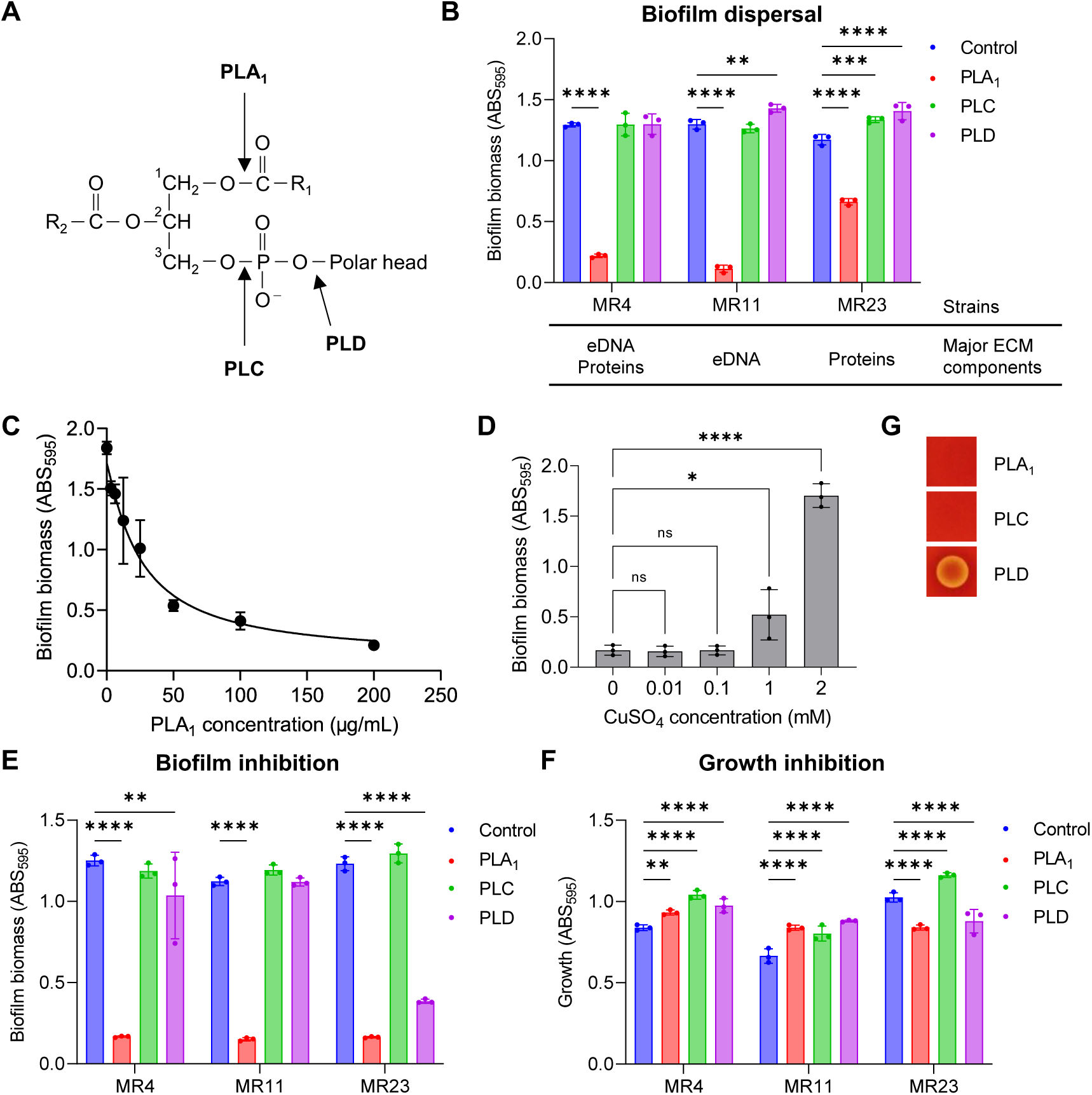
Effects of Phospholipases on *S. aureus* Biofilms. (**A**) Schematic representation of phospholipid digestion sites targeted by the phospholipases used in this study. R1 and R2 represent fatty acid chains, while 1, 2, and 3 denote the *sn* positions of phospholipids. (**B**) Biofilm dispersal analysis. Mature biofilms of the indicated *S. aureus* strains, cultivated in BHIG for 24 h, were treated with the specified bacterial phospholipases or left untreated (Control). Major biofilm components of each strain were previously identified (*9, 10*). Biofilms were stained with crystal violet (CV), and biomass was quantified by measuring absorbance at 595 nm (ABS_595_). (**C**) Dose-dependent effect of PLA_1_ on biofilm degradation. Twenty-four-h biofilms of *S. aureus* strain MR4 were treated with varying concentrations of PLA_1_ at 37°C for 2 h. (**D**) Impact of CuSO_4_, an inhibitor, on biofilm dispersal by 200 μg/ml PLA_1_. (**E**) Inhibition of biofilm formation. The effects of the enzymes on biofilm formation of the indicated strains were evaluated using the CV-staining method. Control samples were cultured without enzymes. (**F**) Effects of the indicated enzymes on *S. aureus* growth. PLA_1_, PLC, and PLD were used at 200 μg/ml, 200 μg/ml, and 1,000 U/ml, respectively, in panels **B**, **E**, and **F**. (**G**) Hemolytic activity of the enzymes was assessed using blood agar plates. Means and standard deviations from at least three independent experiments are presented. Statistical analyses were conducted using two-way ANOVA with Dunnett’s correction for multiple comparisons relative to Control in panels **B**, **E**, and **F**, and one-way ANOVA with Dunnett’s correction for multiple comparisons relative to Control in panel **D**. Statistical significance: *, *p* < 0.05; **, *p* < 0.01; ***, *p* < 0.001; ****, *p* < 0.0001; ns, not significant.

Among the enzymes tested, only bacterial PLA_1_ effectively dispersed pre-formed biofilms of the *S. aureus* strains MR4, MR11, and MR23, which rely on eDNA and/or proteins for biofilm stability (*8–10*) (Fig. 2B). PLA_1_ dispersed the pre-formed biofilm of MR4 in a dose-dependent manner (Fig. 2C), and its biofilm dispersal activity was significantly inhibited by 2 mM CuSO_4_, which inhibits PLA_1_ by direct interaction with the enzyme (*27*) (Fig. 2D). These results indicate the importance of PLA_1_ enzymatic activity for biofilm dispersal.

In addition to dispersing mature biofilms, PLA_1_ also inhibited biofilm formation by the *S. aureus* strains MR4 and MR11 without inhibiting bacterial growth (Fig. 2E-F). Although slight promotion in MR4 and MR11 and inhibition in MR23 by PLA_1_ were observed, these effects were minimal compared to the remarkable inhibition of biofilm formation. Similarly, PLA_1_ prevented biofilm formation in the commonly studied *S. aureus* strains SH1000 and Newman (*28, 29*) without affecting their growth and efficiently dispersed their mature biofilms (fig. S1). This suggests that the biofilm-disrupting activity of PLA_1_ is not limited to specific clinical strains.

Although bacterial PLD did not show biofilm dispersal activity against the MR23 biofilm (Fig. 2B), it was effective at inhibiting biofilm formation when applied from the start (Fig. 2E). For biofilm control applications, enzyme cytotoxicity is a key concern. Notably, none of the tested enzymes, except PLD, exhibited hemolytic activity (Fig. 2G), indicating that PLA_1_ may be safely used in biotechnological and other practical settings.

### PLA_1_ Disrupts Extracellular Nanostructures in Biofilms

To investigate the mechanism behind the anti-biofilm activity of PLA_1_, we observed *S. aureus* biofilms and membranous extracellular nanostructures before and after PLA_1_ treatment. CLSM showed that untreated MR4 biofilms were dense and thick (Fig. 3A), but after PLA_1_ treatment, only a thin layer of biofilm remained on the glass surface (Fig. 3A). Field Emission-Scanning Electron Microscope (FE-SEM), enabling high resolution imaging, showed nanospheres located at the interfaces between *S. aureus* cells in the biofilm, which were absent following PLA_1_ treatment (Fig. 3B).

**Fig. 3.**
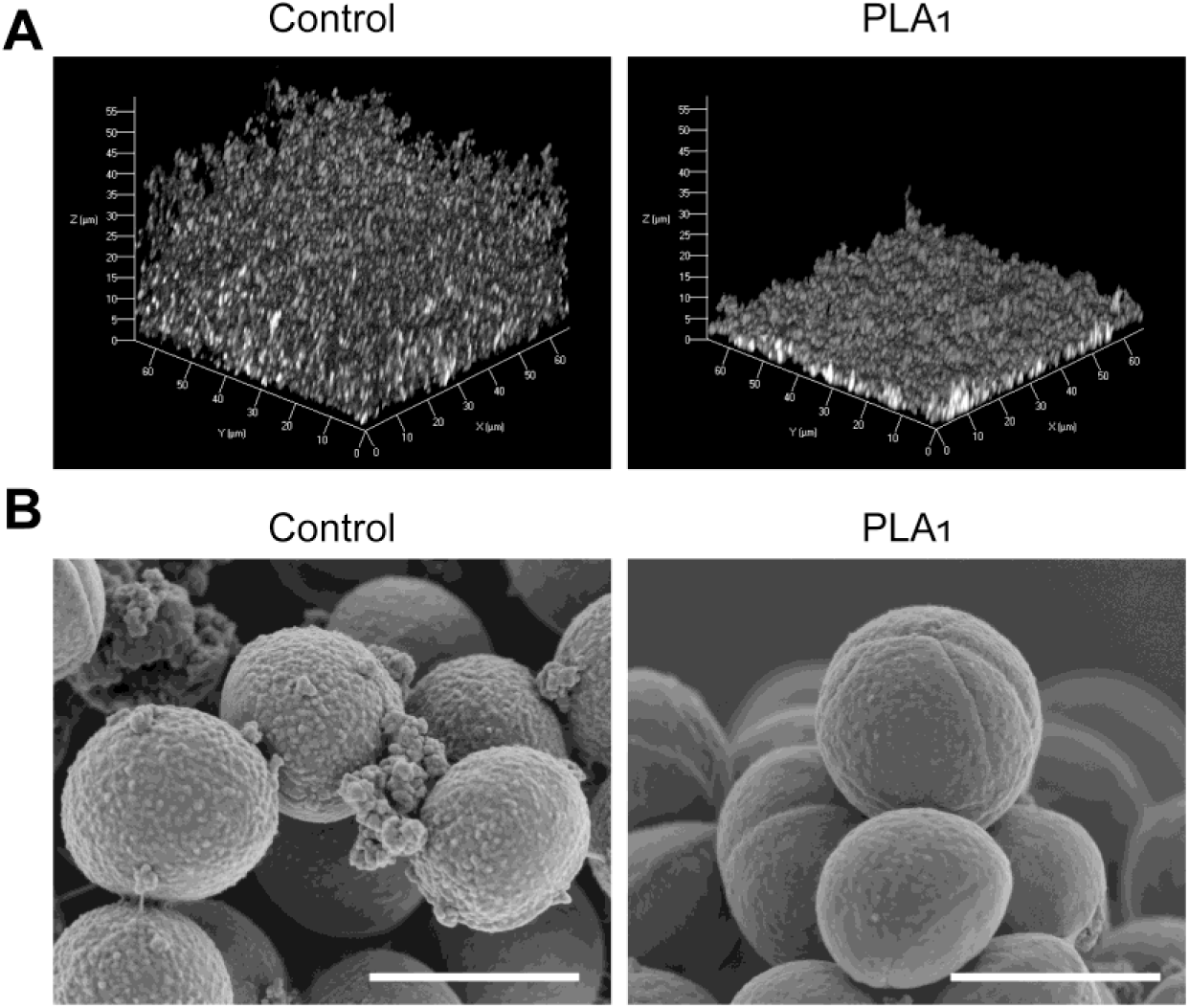
Microscopic Analysis of Biofilms Formed With or Without PLA_1_. Biofilms of *S. aureus* strain MR4 were cultivated in BHIG for 24 h at 37°C, either in the absence (Control) or presence of 200 μg/ml PLA_1_. (**A**) Biofilms formed on glass-bottomed dishes were stained with the membrane-binding probe FM1-43, optically cleared using the iCBiofilm method (*23*), and visualized using CLSM. (**B**) High-resolution SEM images of biofilm cells revealed that PLA_1_ disrupts nanospheres attached to bacterial cell surfaces and at the interface between cells. Scale bars: 1 μm.

Collectively, these findings suggest that PLA_1_ targets extracellular phospholipids in the biofilm matrix, disrupting biofilm integrity and leading to both inhibition of biofilm formation and dispersal of pre-formed biofilms.

### Presence of Phospholipids in the Matrix of *S. aureus* Biofilms

The presence and role of phospholipids in the matrix of *S. aureus* biofilms had not been clearly understood. To address this, we isolated the biofilm matrix from the MR4 strain using a Tris-HCl buffer (pH 8.0) containing 1 M NaCl, following an established protocol (Fig. 4A) (*8*). Thin-layer chromatography (TLC) analysis of the isolated matrix revealed three major phospholipids: phosphatidylglycerol (PG), lysyl-phosphatidylglycerol (Lys-PG), and cardiolipin (CL), all of which are known components of the *S. aureus* membrane (Fig. 4B and fig. S2) (*15, 16*). Notably, these phospholipids disappeared after treatment with PLA_1_, suggesting that they contribute to the structure and stability of the biofilm and are susceptible to enzymatic cleavage, potentially facilitating biofilm dispersion and inhibition.

**Fig. 4.**
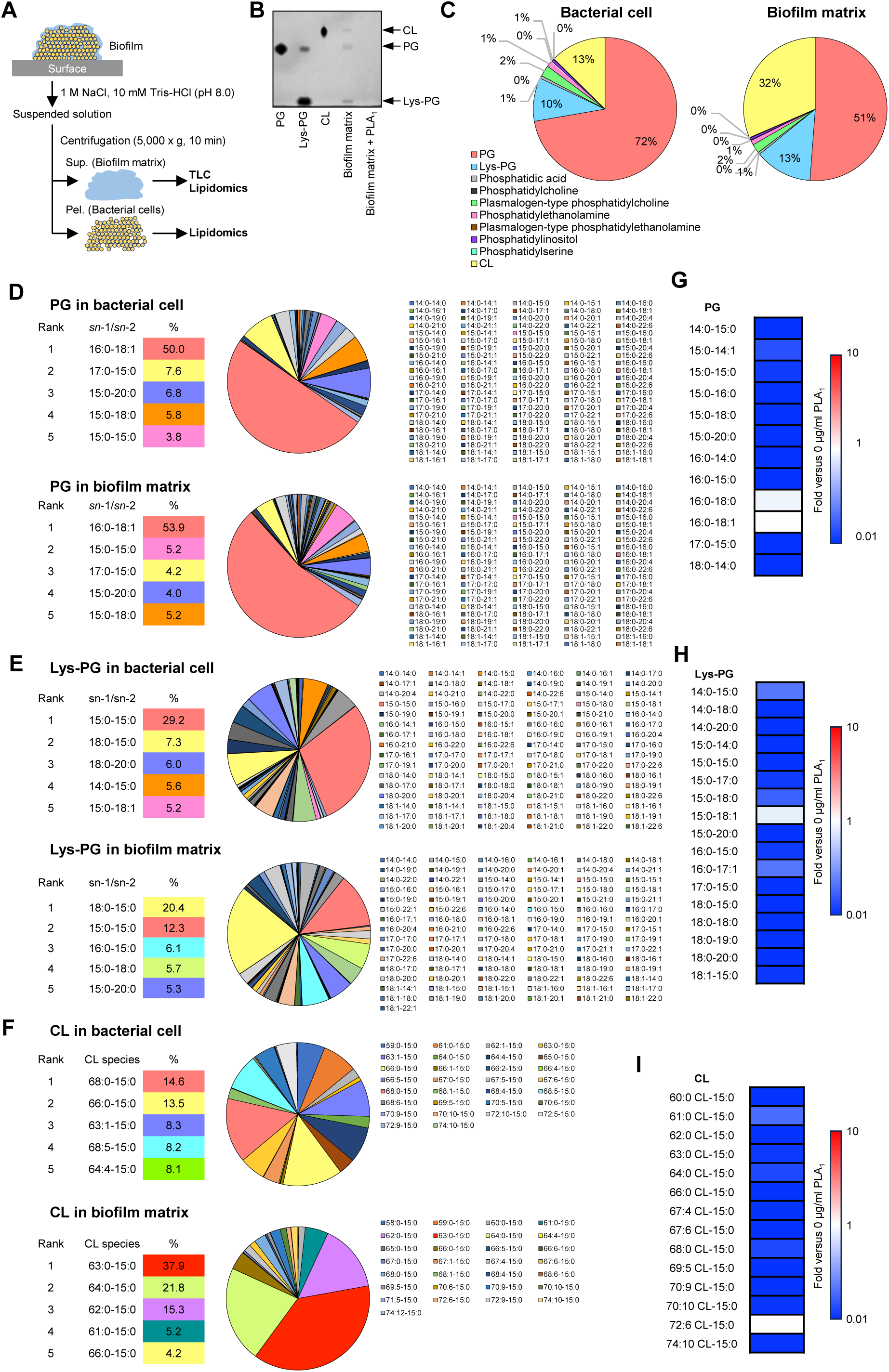
Phospholipid Composition Analysis via Lipidomics. (**A**) Schematic representation of TLC and lipidomic analyses for assessing phospholipid composition in the biofilm matrix and bacterial cells from the 24-h MR4 biofilm. (**B**) Phospholipid contents were analyzed using TLC, with purified Lys-PG (C_16:0_), PG (C_16:0_), and CL (C_16:0_) as standards. (**C**) Phospholipid composition in *S. aureus* MR4 cells and the biofilm matrix, determined by lipidomic analysis as outlined in the Methods. (**D-F**) Fatty acid acyl chain composition of PG (**D**), Lys-PG (**E**), and CL (**F**) in MR4 cells and the biofilm matrix. (**G-I**) Relative levels of major PG (**G**), Lys-PG (**H**), and CL (**I**) species after treatment with 20 μg/ml PLA_1_, expressed as ratios to pre-treatment levels (MS intensity > 10^6^ for PG and CL, > 10^5^ for Lys-PG) in the biofilm matrix of MR4. The abundance of each phospholipid species prior to PLA_1_ treatment is normalized to 1.

To further characterize the phospholipid composition in the biofilm matrix, we conducted a detailed lipidomic analysis. Comparisons between the biofilm matrix and *S. aureus* cells revealed that PG was the most abundant phospholipid in both contexts (Fig. 4C). However, CL and Lys-PG were slightly enriched in the biofilm matrix, increasing from 13% and 10% in bacterial cells to 32% and 13%, respectively (Fig. 4C).

We then analyzed the fatty acid acyl chain profiles of PG, Lys-PG, and CL. In bacterial cells, the predominant PG species was C_16:0-18:1_ (50.0%), while the most abundant Lys-PG species was C_15:0-15:0_ (29.2%) (Fig. 4, D and E). In the biofilm matrix, Lys-PG species such as C_18:0-15:0_ (20.4%), C_16:0-15:0_ (6.1%), and C_15:0-18:0_ (5.7%) were enriched (Fig. 4E). In contrast, PG species showed no obvious enrichment in the matrix (Fig. 4D). For CL, we focused on species containing C_15:0_, a major fatty acid in PG and Lys-PG, due to analytical limitations. Enrichment of specific CL species, including CL (C_63:0-15:0_), CL (C_64:0-15:0_), CL (C_62:0-15:0_), and CL (C_61:0-15:0_), was observed in the biofilm matrix (Fig. 4F).

Finally, we investigated the effects of PLA_1_ treatment on the fatty acid acyl chains of these phospholipids in the isolated biofilm matrix. PLA_1_ effectively cleaved various PG, Lys-PG, and CL species, regardless of their fatty acid composition, demonstrating broad activity against phospholipid substrates (Fig. 4, G–I, and fig. S3).

In summary, our lipidomic analysis highlights an enrichment of specific Lys-PG and CL species in the *S. aureus* biofilm matrix. These phospholipids are susceptible to enzymatic degradation by PLA_1_, offering potential insights into mechanisms for biofilm disruption and inhibition.

### Lys-PG Induces *S. aureus* Cell Aggregation Through Electrostatic and Hydrophobic Interactions

Given the presence of nanospheres at the interfaces between *S. aureus* cells within biofilms (Fig. 3B), we hypothesized that extracellular phospholipids may act as molecular glues, linking bacterial cells. To explore this, we tested the effects of the three major phospholipids identified in our TLC and lipidomic analyses on *S. aureus* cell aggregation. In plastic tubes, Lys-PG (C_18:1_) induced cell aggregation in a dose-dependent manner, whereas PG (C_18:1_) and CL (C_18:1_) produced less aggregates (Fig. 5A).

**Fig. 5.**
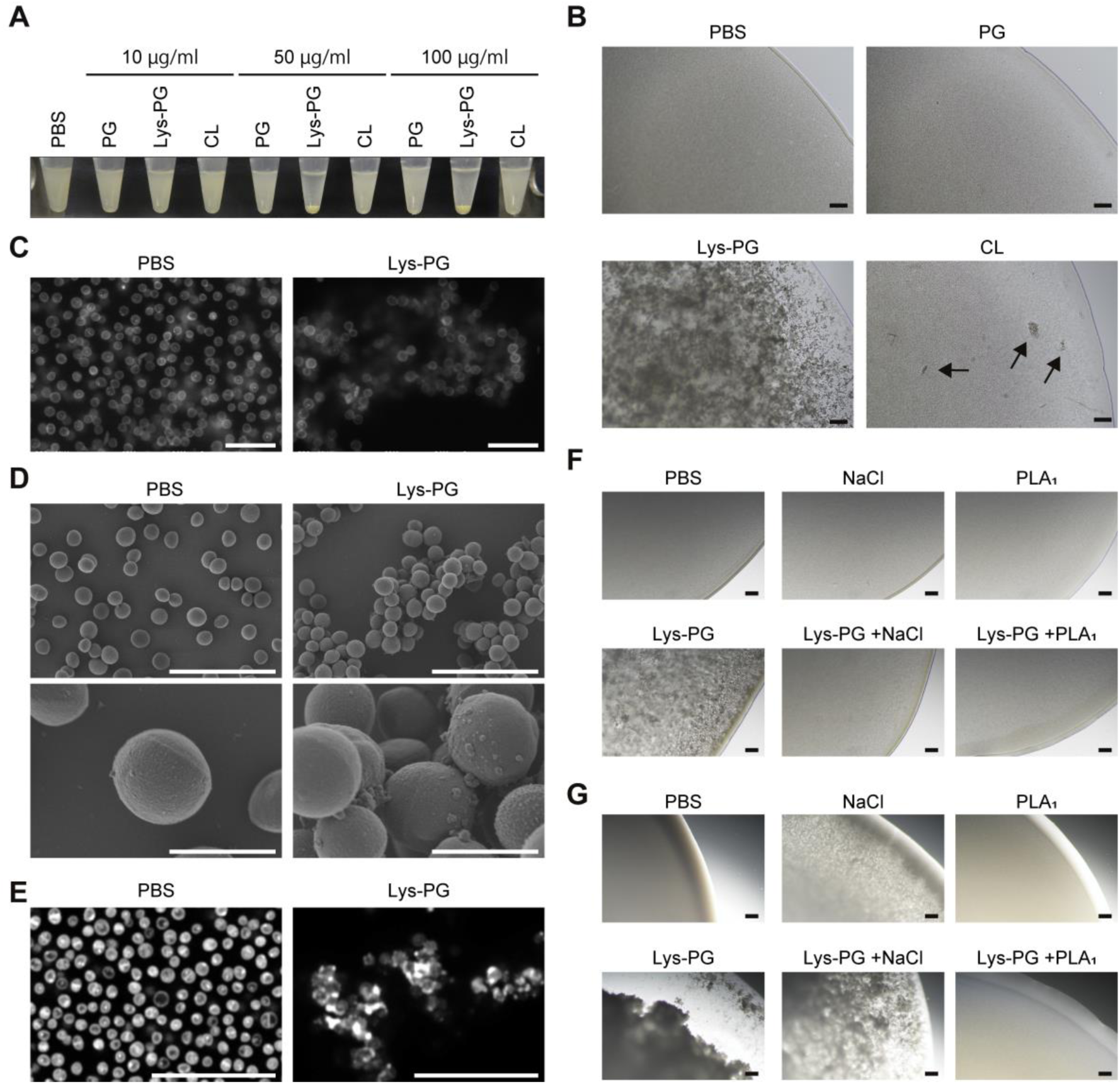
Impact of Phospholipids on *S. aureus* Cell Aggregation. (**A**) The effects of purified phospholipids on *S. aureus* MR4 cell aggregation were examined in 1.5 ml plastic tubes. After incubation in PBS supplemented with the indicated concentrations of Lys-PG (C_18:1_), PG (C_18:1_), or CL (C_18:1_) at 25°C for 30 min, photos of the tubes were taken. (**B**) *S. aureus* MR4 cells were mixed with 100 μg/ml PG (C_18:1_), 100 μg/ml Lys-PG (C_18:1_), or 100 μg/ml CL (C_18:1_) in PBS, and then, small aliquots (10 μl) were placed on a slide glass. After incubation at 25°C for 30 min, cell aggregation of *S. aureus* was directly observed by optical microscopy with a 10× objective lens. Arrows indicate small aggregates. (**C**) *S. aureus* MR4 cells were incubated in the presence or absence of 100 μg/ml Lys-PG (C_18:1_) in PBS on ASEM dishes. After incubation at 25°C for 30 min, cells were fixed with 1% glutaraldehyde, stained with positively charged nanogold and phosphotungstic acid, and observed by ASEM. (**D, E**) *S. aureus* MR4 cells were incubated in the presence of 100 μg/ml Lys-PG (C_18:1_) and FM1-43 in PBS in a plastic tube. After incubation at 25°C for 30 min, cells were observed by SEM (**D**) or CLSM with 63× objective lens and an Airyscan high resolution unit (**E**). (**F**) *S. aureus* MR4 cells were mixed with 100 μg/ml Lys-PG (C_18:1_), 1 M NaCl, or 200 μg/ml PLA_1_ in 1.5 ml plastic tubes. If required, 1 M NaCl and 200 μg/ml PLA_1_ were used with 100 μg/ml Lys-PG (C_18:1_). After incubation at 25°C for 30 min, photos of cell aggregates were taken by optical microscopy with 10× objective lens. (**G**) Negatively charged microbeads with a diameter of 0.9 μm were mixed with 100 μg/ml Lys-PG (C_18:1_), 1 M NaCl, 200 μg/ml PLA_1_, 100 μg/ml Lys-PG (C_18:1_) and 1 M NaCl, or 100 μg/ml Lys-PG (C_18:1_) and 200 μg/ml PLA_1_ in 1.5 ml plastic tubes. After incubation at 25°C for 30 min, aggregates of beads were observed by microscopy with a 10× objective lens. As a control, phospholipids were not added. Representative images from three independent experiments are shown. Scales are 1 μm in the lower panels of **D**, 5 μm in **C**, 10 μm in the upper panels of **D** and **E**, and 100 μm in **B**, **F**, and **G**.

We next examined cell aggregation on a microscale using bright-field microscopy with a 10× objective lens. Lys-PG (C_18:1_) triggered the rapid formation of large aggregates within 30 min (Fig. 5B). In contrast, PG (C_18:1_) did not cause noticeable aggregation, and CL (C_18:1_) had only a minimal effect (Fig. 5B). Similar aggregation patterns were observed with phospholipids composed of different fatty acid chain lengths and saturation levels, such as C_16:0_ (fig. S4). Real-time light scattering measurements of bacterial cell suspensions revealed a slight decrease within the first 5 min, followed by a sharp decline after 10–15 min upon the addition of Lys-PG (C_18:1_) (fig. S5). These results suggest that Lys-PG (C_18:1_) induces *S. aureus* aggregation through two distinct phases: an initial small aggregate formation phase (∼5 min) followed by the formation of larger aggregates (10– 15 min).

Among the phospholipids tested, Lys-PG (C_18:1_) had the most pronounced impact on cell aggregation. To further investigate this, we used ASEM to visualize cell aggregates and surface attachment in the presence of Lys-PG (C_18:1_) under aqueous conditions. While separated and small aggregated cells of *S. aureus* attached to the ASEM dish, they formed extensive aggregates when exposed to Lys-PG (C_18:1_) (Fig. 5C and fig. S6A), consistent with the optical microscopy observations. FE-SEM also showed that Lys-PG (C_18:1_) induced aggregation of *S. aureus* cells, with nano-spherical structures visible at the interfaces between the cells (Fig. 5D).

To confirm Lys-PG localization within aggregates of *S. aureus* cells, the cells treated with Lys-PG (C_18:1_) were stained with FM1-43 and observed by CLSM in super-resolution mode. In the absence of Lys-PG, monococci and diplococci labeled with FM1-43 adhered to the surface (Fig. 5E and fig. S6B). In contrast, Lys-PG (C_18:1_)-treated cells formed large aggregates with strong fluorescence, typically localized between cells (Fig. 5E and fig. S6B), indicating that Lys-PG (C_18:1_) is concentrated at cell interfaces, likely acting as a molecular glue that connects *S. aureus* cells in three dimensions.

Since Lys-PG, but not PG, strongly promoted cell aggregation, we hypothesized that the positively charged lysine moiety of Lys-PG (fig. S2) mediates aggregation through electrostatic interactions with the negatively charged bacterial surface. To test this, we used a high-concentration salt solution (1 M NaCl) to disrupt electrostatic interactions, which successfully prevented Lys-PG (C_18:1_)-induced bacterial cell aggregation (Fig. 5F). In addition, treatment with PLA_1_, which cleaves the ester bonds in Lys-PG, also suppressed aggregation (Fig. 5F), suggesting that the hydrophobic moiety of Lys-PG contributes to aggregation through hydrophobic interactions between Lys-PG molecules.

Next, we explored whether specific surface molecules on *S. aureus* are necessary for Lys-PG-induced aggregation. In *S. aureus*, several negatively charged molecules on the cell surface are known such as teichoic acids and cell wall-anchored proteins. We tested a wall teichoic acid-deficient (Δ*tagO*) strain of *S. aureus* and proteinase K-treated wild-type cells to degrade surface proteins. Both Δ*tagO* and proteinase K-treated wild-type cells formed large aggregates within 30 min, similar to intact wild-type cells. However, proteinase K-treated Δ*tagO* cells only formed smaller aggregates compared with the other cases (fig. S7), indicating that negatively charged molecules play a role in Lys-PG-triggered aggregation.

We then tested whether the negative charge alone is sufficient for Lys-PG-triggered aggregation by using silica microbeads (0.9 μm diameter) with a negatively charged surface, mimicking *S. aureus* cells. Lys-PG (C_18:1_) strongly induced aggregation of microbeads, which was suppressed by PLA_1_ and partially inhibited by high concentrations of NaCl (Fig. 5G).

Collectively, these findings demonstrate that Lys-PG functions as a molecular glue mediating cell-to-cell adhesion during biofilm formation. Both electrostatic interactions between the positively charged Lys-PG and the negatively charged bacterial surface, and hydrophobic interactions between fatty acid acyl chains of Lys-PG molecules, are critical for Lys-PG-induced aggregation (Fig. 6). Importantly, no specific surface molecules are required for this under the conditions tested.

**Fig. 6.**
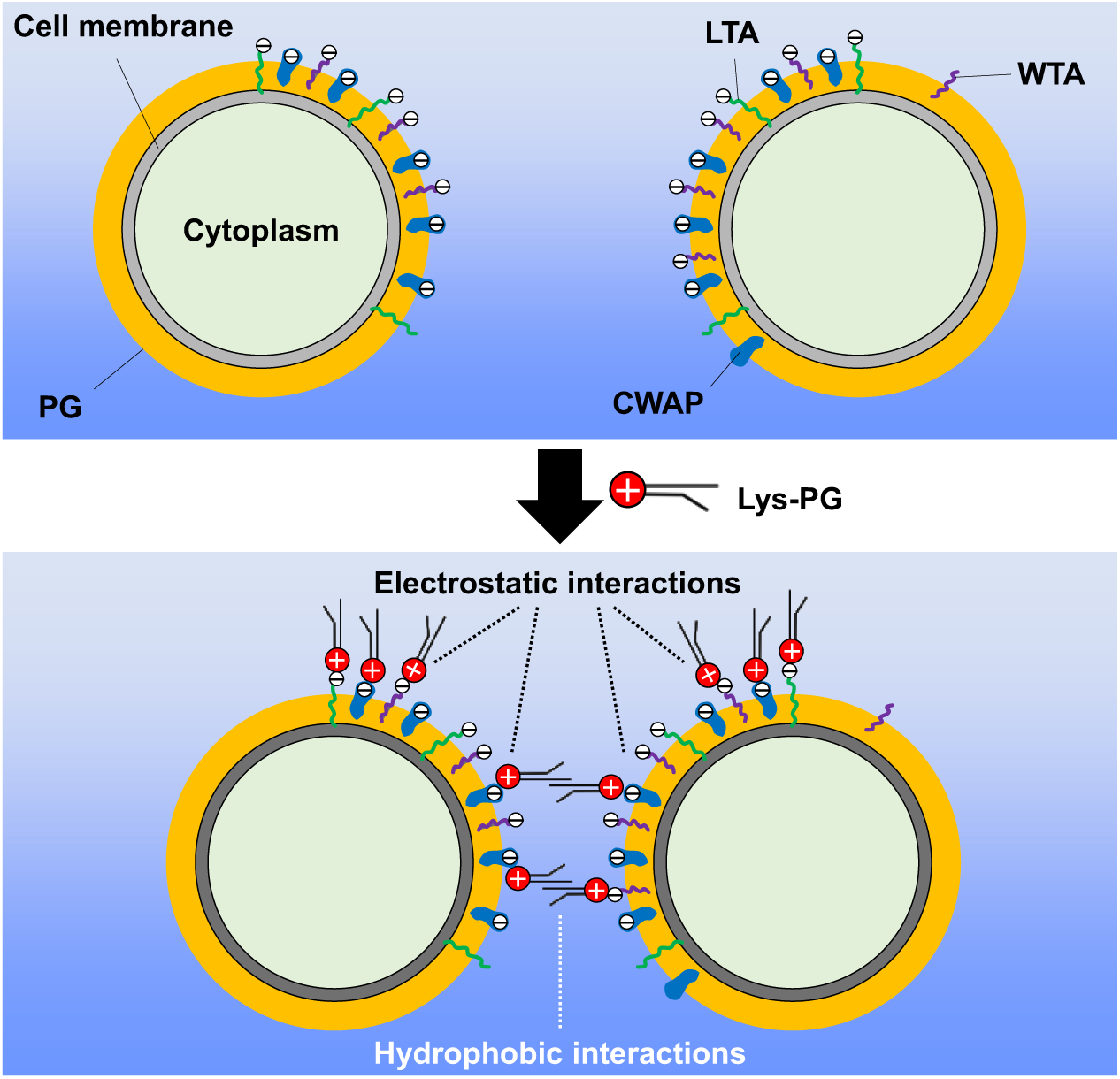
Proposed Model for the Role of Lys-PG in Cell Aggregation. The positively charged lysine (Lys) moiety of Lys-PG can bind to negatively charged molecules on the bacterial surface, such as wall teichoic acids (WTA), lipoteichoic acids (LTA), and cell wall-anchored proteins (CWAP), via electrostatic interactions. Additionally, at least two Lys-PG molecules can interact with each other through their hydrophobic fatty acid tails. Therefore, both the electrostatic interactions between the Lys moieties of Lys-PG and the bacterial cell surface, as well as the hydrophobic interactions between Lys-PG molecules, play important roles in mediating cell aggregation.

### Lys-PG Is Crucial for Biofilm Formation in *S. aureus*

To assess whether Lys-PG is involved in biofilm formation in *S. aureus*, we generated an *S. aureus* MR4 mutant strain in which the gene encoding the Lys-PG synthesis enzyme (*mprF*) was deleted. TLC analysis confirmed that the MR4 Δ*mprF* mutant lacked Lys-PG, while the production of phosphatidylglycerol (PG) and cardiolipin (CL) remained unaffected (Fig. 7A), in agreement with previous studies (*30, 31*).

**Fig. 7.**
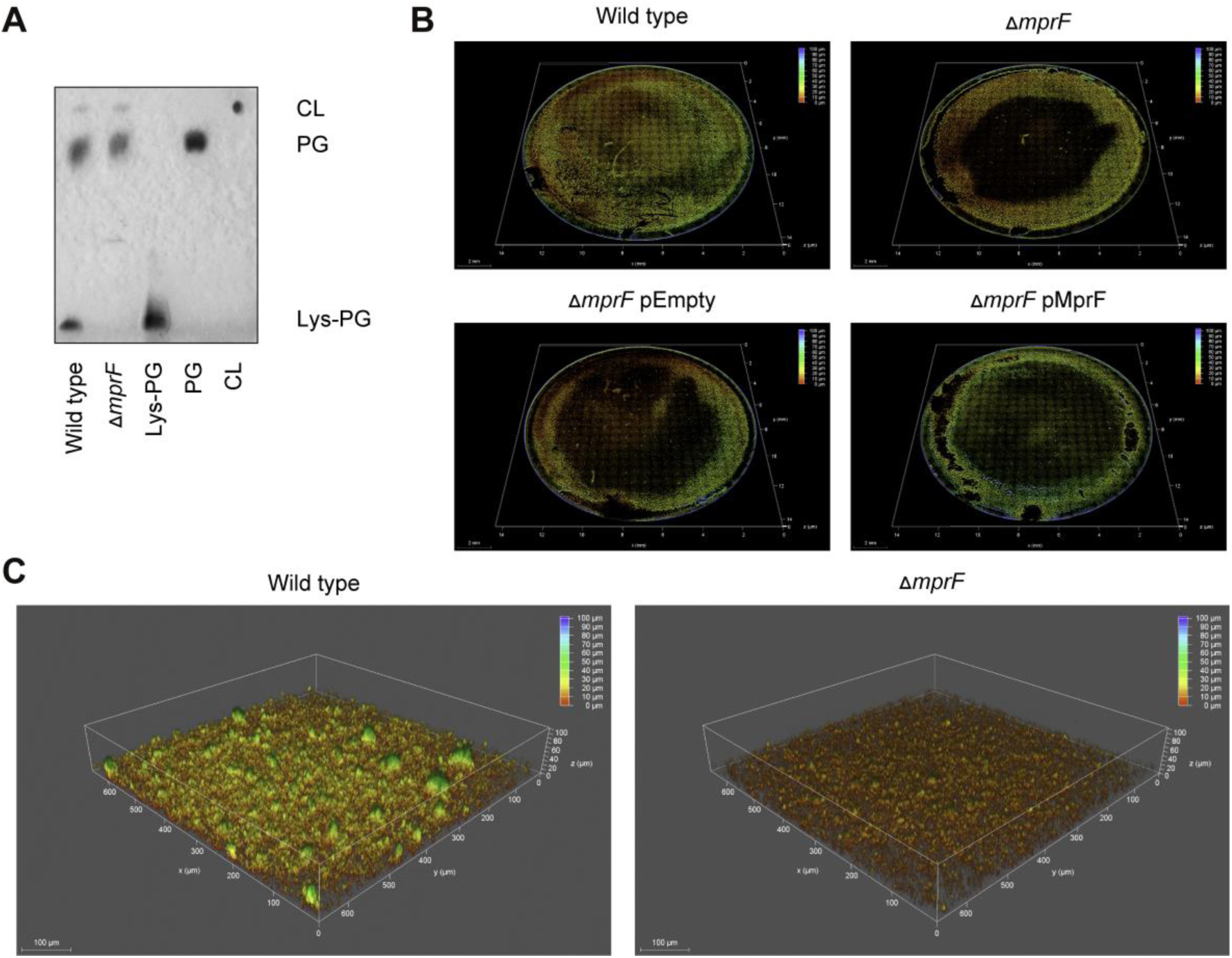
Effect of Defective Lys-PG Production on Biofilm Formation. (**A**) Membrane phospholipids in the MR4 wild-type and Δ*mprF* strains were analyzed by TLC. (**B** and **C**) Biofilms formed by the indicated strains were stained with FM1-43 and observed on the THUNDER Imaging system with a 20× objective (Leica). (**B**) Wide-field views are presented to show non-biased images of entire biofilms formed on the glass surface. (**C**) Representative enlarged images of the biofilms formed by the MR4 wild-type and Δ*mprF* strains.

We then compared the biofilm formation of MR4 Δ*mprF* with the wild-type strain. To ensure accurate and comprehensive observation, biofilms were visualized using the Thunder Imaging System (*23*) with tiling mode to capture the entire biofilm area on glass-bottom dishes. The MR4 Δ*mprF* strain formed a significantly thinner biofilm compared to the wild-type strain (Fig. 7B), a result consistent with previous findings using the MRSA USA300 strain JE2 and its *mprF* mutant (*32*). Notably, the central region of the Δ*mprF* biofilm was particularly reduced in thickness compared to the wild type (Fig. 7, B and C). Reintroducing the *mprF* gene via a plasmid partially restored biofilm formation in the mutant (Fig. 7B).

These results demonstrate the critical role of Lys-PG in *S. aureus* biofilm formation, especially in maintaining biofilm thickness, highlighting its importance in promoting a robust biofilm structure.

## DISCUSSION

In this study, we investigated the role of extracellular phospholipids in *S. aureus* biofilms, particularly focusing on the moonlighting function of Lys-PG in promoting bacterial aggregation and biofilm formation. While extracellular nucleic acids (eDNA and envRNA), proteins, and polysaccharides have long been recognized as key structural components of *S. aureus* biofilms (*10, 12*), this study demonstrates the previously overlooked importance of extracellular phospholipids. Notably, the discovery that Lys-PG is enriched in the biofilm matrix and promotes cell aggregation adds a novel layer to our understanding of biofilm structure. Role of Lys-PG as a molecular glue within *S. aureus* biofilms offers new insights into biofilm architecture, where phospholipids—once thought to play a minor or undefined role in the biofilm matrix—are now seen as pivotal in forming intercellular junctions. Our results also demonstrate how the enzymatic activity of PLA_1_ can inhibit biofilm development and disperse pre-formed biofilms.

The selective enrichment of specific Lys-PG and CL species, such as Lys-PG (C_18:0-15:0_, C_16:0-15:0_, and C_15:0-18:0_) and CL (C_63:0-15:0_, C_64:0-15:0_, and C_62:0-15:0_), within the biofilm matrix compared to their relative abundance in bacterial cells (Fig. 4, E and F) suggests a specialized role for these phospholipids in maintaining biofilm structural integrity. These enriched phospholipid species may interact with other components of the biofilm matrix, such as extracellular DNA (eDNA), proteins, and polysaccharides, or with structural elements on bacterial cells, including cell wall peptidoglycan and teichoic acids. Unraveling the molecular mechanisms underlying the selective accumulation of these phospholipids in the biofilm matrix should be a key area of focus for future studies. Our study demonstrates that Lys-PG triggers bacterial cell aggregation via electrostatic and hydrophobic interactions, with the positively charged lysine moiety mediating electrostatic adhesion to negatively charged bacterial surfaces. This mechanism is supported by experiments showing that high concentrations of NaCl can disrupt these interactions, and PLA_1_ can cleave Lys-PG to prevent aggregation. Since our lipidomic analysis did not detect lyso-Lys-PG in the PLA_1_-treated samples, it is likely that PLA_1_ cleaved Lys-PG at both the *sn*-1 and *sn*-2 positions under the tested conditions. The dual role of Lys-PG—combining electrostatic and hydrophobic forces—provides a robust and flexible mechanism for biofilm stability. Additionally, we observed that CL also indued *S. aureus* cell aggregation in vitro (Fig. 5B and fig. S4). Unlike Lys-PG, CL lacks a positively charged lysine moiety, suggesting that CL may act through a distinct mechanism.

PLA_1_ stands out as a promising biofilm dispersal agent in this study, primarily due to its ability to cleave Lys-PG and other phospholipids in the biofilm matrix without inhibiting bacterial growth. This selective activity is particularly valuable for both clinical and industrial applications, where removing biofilms without killing bacteria may be crucial, such as in scenarios requiring the preservation of microbial viability for ecosystem balance or fermentation processes. The limited capacity of PLA_1_ to inhibit bacterial growth, as demonstrated in Fig. 2E and fig. S1, may result from the robust cell wall of *S. aureus*, which likely limits enzyme access to the bacterial membrane, or from compensatory mechanisms that facilitate de novo phospholipid synthesis in response to PLA_1_ activity (*33*). Moreover, the absence of hemolytic activity associated with PLA_1_ (Fig. 2G) further supports its potential for safe application in various biotechnological contexts. This safety profile is particularly significant for use in medical device coatings or surfaces that come into contact with human tissues, where minimizing cytotoxicity is a critical consideration.

This report raises several important questions for future research. For instance, while Lys-PG is shown to play an important role in *S. aureus* biofilm formation, it remains unclear whether similar phospholipid-based mechanisms operate in other pathogenic bacteria. Comparative studies on phospholipid content in biofilms formed by different bacterial species could reveal whether PLA_1_ or related enzymes could be universally applied to biofilm control. Additionally, the report highlights the need for further exploration into the structure-function relationships of different Lys-PG species. The enrichment of specific Lys-PG species in the biofilm matrix suggests that particular acyl chain compositions may be more effective at promoting biofilm integrity. Investigating these differences could lead to the development of tailored biofilm dispersal agents or coatings that specifically disrupt biofilms under different environmental conditions. Lastly, the report hints at the potential roles of phospholipases in both in vivo and environmental settings. Host-derived phospholipases may influence the survival and colonization of beneficial or pathogenic bacteria by inhibiting or dispersing biofilms. Similarly, microbe-derived phospholipases could be utilized to selectively disrupt biofilms in environmental niches with mixed bacterial communities. These insights open promising avenues for employing phospholipases as targeted tools in biofilm management and microbial ecosystem regulation.

## Conclusion

This study provides groundbreaking insights into the role of extracellular phospholipids in *S. aureus* biofilm formation and highlights the potential of PLA_1_ as an effective biofilm dispersal agent. By uncovering the moonlighting function of Lys-PG, this research broadens our understanding of biofilm architecture and resilience while opening new avenues for the development of innovative biofilm control strategies in both clinical and industrial settings. These findings could inspire further investigations into phospholipase-based treatments and their applications in biofilm management.

## METHODS

### Bacterial strains and media

Bacterial strains used in this study are listed in Table S1. Clinical strains of *S. aureus* (*10*) were isolated from patients in The Jikei University Hospital (Table S1). These strains were grown in brain heart infusion (BHI) (Becton Dickinson, Franklin Lakes, NJ, USA). For cultivation of *E. coli*, Lysogeny Broth (LB) (1% tryptone, 0.5 % yeast extract, and 0.5% NaCl) was used. For selection of transformants, appropriate antibiotics (100 μg/ml ampicillin and 5 μg/ml chloramphenicol; Nacalai Tesque, Kyoto, Japan) and an inducer (100 ng/ml aTc; Sigma, St. Louis, MO, USA) were added to the media.

### Construction of knockout strains

The 500-bp upstream and downstream sequences of *mprF* from MR4 genomic DNA were amplified using the polymerase chain reaction (PCR) with the following primer sets: attB1-mprF-F; mprF-R, and mprF-F; attB2-mprF-R (Table S2). These fragments were ligated by splicing using an overlap extension PCR (*9, 11, 12*). The resulting fragment was cloned into pKOR1 using Gateway BP Clonase II enzyme mix (Thermo Fisher Scientific, Waltham, MA, USA). The generated plasmid was named pMprF-KO (Table S1). Using pMprF-KO, *mprF* was deleted from MR4 by in-frame deletion according to a previously reported procedure (*11, 12, 34*). Briefly, pMprF-KO was used to transform *S. aureus* RN4220 (*35*) by electroporation. After purification from RN4220, the plasmid was introduced into MR4 by electroporation. The transformant harboring pMprF-KO was cultured on NYE agar (1% casein hydrolysate, 1% yeast extract, 0.5% NaCl, pH 7.2) containing 5 μg/ml chloramphenicol at 30°C for 24 h. A single colony on the plate was inoculated into BHI containing 5 μg/ml chloramphenicol and grown at 30°C for 48 h. The culture was diluted 1,000-fold into BHI containing 5 μg/ml chloramphenicol and incubated at 42°C overnight. The overnight culture (50 μl) was spread on tryptic soy broth (TSB, Becton Dickinson) agar containing 20 μg/ml chloramphenicol and incubated at 42°C overnight. A single colony was inoculated into TSB and grown at 30°C overnight. The overnight culture was diluted 100-fold into TSB, and the aliquot (100 μl) was spread on TSB agar containing 100 ng/ml anhydrotetracycline. After incubation at 30°C overnight, several big colonies on the plate were selected as the MR4 Δ*mprF* candidates and grown in TSB at 37°C overnight. Deletion of *mprF* was confirmed by PCR.

For complementation of *mprF*, the strain MR4 Δ*mprF* was transformed by electroporation using pHYmrpF (*31*). As a control, the same strain was transformed by electroporation using pHY300PLK (Takara, Shiga, Japan).

### Enzymes

Proteinase K from *Tritirachium album*, PLA_1_ from *Thermomyces lanuginosus* (≥10 KLU/g liquid), PLC from *Bacillus cereus* (≥1,000 units/mg protein), and PLD from *Streptomyces chromofuscus* (≥50,000 U/ml) were procured from Sigma-Aldrich.

### Phospholipids

1,2-dioleoyl-*sn*-glycero-3-[phospho-rac-(3-lysyl(1-glycerol))] (Lys-PG with C_18:1_), 1,2-dipalmitoyl-*sn*-glycero-3-[phospho-rac-(3-lysyl(1-glycerol))] (chloride salt) (Lys-PG with C_16:0_), 1,2-dioleoyl-*sn*-glycero-3-phospho-(1’-rac-glycerol) (sodium salt) (PG with C_18:1_), 1,2-dipalmitoyl-*sn*-glycero-3-phospho-(1’-rac-glycerol) (sodium salt) (PG with C_16:0_), 1’,3’-bis[1,2-dioleoyl-*sn*-glycero-3-phospho]-glycerol (sodium salt) (CL with C_18:1_), and 1,2-dipalmitoyl-*sn*-glycero-3-ethylphosphocholine (chloride salt) (CL with C_16:0_) were purchased from Avanti Polar Lipids (Birmingham, AL, USA). The lyophilized powders were dissolved in 100% methanol, 100% chloroform, or chloroform/methanol (9 : 1, v/v).

### Bacterial growth monitoring

Growth of the indicated strains was monitored at 37°C in BHIG in the absence and presence of PLA_1_ in 96-well flat bottom polystyrene plates (Corning, NY, USA) by measuring the absorbance of each culture at 600 nm every 30 min for 24 h using a Bio Microplate Reader HiTS (Scinics Corp., Tokyo, Japan).

### Biofilm inhibition assay

Overnight cultures grown in BHI medium at 37°C were diluted 1,000-fold in BHI supplemented with 1% (w/v) glucose (BHIG). Two hundred microliters of these suspensions were cultured in 96-well flat bottom polystyrene plates (Corning, NY, USA) at 37°C for 24 h. When required, PLA_1_ (200 μg/ml), PLC (200 μg/ml), or PLD (500 U/ml) were added to culture media at the onset of biofilm formation. These biofilms were then washed twice with 150 μl of phosphate-buffered saline (PBS) and stained with 0.05% (w/v) Crystal Violet (CV) (150 μl) for about 3 min at 25 °C. After staining, biofilms were washed once with PBS and biofilm biomass was quantified by measuring the absorbance at 595 nm with an Infinite F200 Pro (Tecan, Männedorf, Switzerland) microtiter plate reader.

### Biofilm destruction assay

Overnight cultures of *S. aureus* grown in BHI medium at 37°C were diluted 1,000-fold in BHIG. Two hundred microliters of these suspensions were cultured in 96-well flat bottom polystyrene plates (Corning, NY, USA) at 37°C for 24 h. Subsequently, PLA_1_ (200 μg/ml or the indicated concentrations), PLC (200 μg/ml), or PLD (1,000 U/ml) were added directly to 24 h-old biofilms without removal of the culture medium and the mixtures were incubated for 2 h at 37°C. The biofilms were quantified as described above.

### Isolation of biofilm matrixes

Biofilm matrixes were extracted from bacteria grown under biofilm formation conditions as previously reported (*8*). Briefly, 24-h biofilms of MR4 formed in 30 ml BHIG were collected by centrifugation at 5,000 × g for 10 min at 25°C. The pellets were centrifuged again for 1 min and the supernatants were removed completely. Then, the pellets were suspended in 10 mM Tris-HCl buffer (pH 8.0) containing 1 M NaCl solution (100 μl) and mixed by vortex. After that, the suspensions were centrifuged at 5,000 × g for 10 min at 25°C. The supernatants were collected as biofilm matrix fractions.

### TLC analysis

Phospholipids were extracted from biofilm cells and isolated biofilm matrixes for TLC analysis using the following procedure.

Biofilms of *S. aureus* wild-type and Δ*mprF* strains were cultivated in 20 ml of BHIG medium in polystyrene plates at 37°C for 24 h under static conditions. After incubation, the culture supernatant and planktonic cells were removed, and the biofilms were resuspended in 20 ml of PBS. The suspensions were centrifuged at 10,000 ×g for 10 min at 4°C, and the supernatants were discarded. The resulting pellets were washed three times with 20 ml of PBS. The washed pellets were resuspended in 0.12 M sodium acetate buffer (pH 4.8) and mixed with 5 ml of chloroform and 11 ml of methanol. After vigorous agitation at 25°C for 60 min, the samples were centrifuged at 15,000 ×g for 5 min at 20°C. The supernatants were transferred to fresh test tubes, followed by the addition of 5 ml chloroform and 5 ml distilled water. After another round of centrifugation at 15,000 ×g for 5 min at 20°C, the lower organic phase was collected and vacuum-dried using a SpeedVac (Thermo Fisher Scientific) at 45°C for 30 min. The dried phospholipids were then completely dissolved in 0.1 ml of chloroform through sonication and prepared for TLC analysis.

For the biofilm matrixes, 500 μl of the isolated matrix was mixed with 500 μl of an organic solvent mixture (chloroform : methanol = 2 : 1, v/v). After vigorous mixing at 25°C for 30 min, the solution was centrifuged at 15,000 ×g for 5 min at 20°C. The lower organic phase was collected, vacuum-dried using a SpeedVac at 45°C for 30 min, and the phospholipids were fully dissolved in 10 μl of the same organic solvent mixture via sonication for subsequent TLC analysis.

Extracted lipids were separated using TLC on a silica gel plate (SilicaGel 60 with concentrating zone, Millipore, Burlington, MA, USA), developed with a solvent mixture of chloroform/methanol/acetic acid (65 : 25 : 10, v/v/v). For lipid detection, TLC plates were sprayed with molybdenum blue reagent (Sigma). PG (C_16:0_), Lys-PG (C_16:0_), and CL (C_16:0_) were used as standards.

### LC-MS/MS analysis

The isolated biofilm matrix (20 μl) was 5-fold diluted into in BFD buffer [20 mM HEPES-OH (pH 7.4) and 20 mM CaCl_2_] supplemented with or without PLA_1_ (200 μg/ml) at 37°C for 3 h. Then, the reaction was stopped by adding 900 μl methanol. Subsequently, phospholipids were analyzed by QTRAP6500^+^, a triple quadrupole-linear ion trap hybrid mass spectrometer (Sciex, Framingham, MA, USA) equipped with a ExionLC^TM^ AD UPLC system (Sciex, Framingham, MA, USA) as follows. Lipids were extracted from the reactants using the modified Bligh and Dyer method.

For detection of PG, Lys-PG and CL, separation and analysis were performed as reported previously (*36*). A 5 µL of each sample was applied to ACQUITY UPLC HSS C18 column (2.1 x 150 mm, 1.8 µm particle size, Waters Corporation, Milford, MA, USA) coupled to ESI-MS, and separated by a linear gradient with mobile phase A (acetonitrile/water = 3:2 [v/v] containing 0.1% formic acid, and 10 mM ammonium acetate) and mobile phase B (isopropyl alcohol/acetonitrile = 9:1 [v/v] containing 0.1% formic acid, and 10 mM ammonium acetate) at a flow rate of 0.22 mL/min at 40°C.

Quantification of PG and the other phospholipids were performed according to previously published method (*37, 38*). Briefly, 5 µL of each sample was separated with a Kinetex C18 column (1.7 μm, 150 × 2.1 mm, Phenomenex) maintained at 50°C, using mobile phase A (acetonitrile/methanol/water = 1:1:1 [v/v/v] containing 0.1% formic acid and 1 mM ammonium formate) and mobile phase B (2-propanol containing 0.1% formic acid and 1 mM ammonium formate) in a linear gradient program with a flow rate of 0.2 ml/min.

Phospholipids were identified based on multiple reaction monitoring (MRM) transitions and retention times. As internal standards, 16:0-*d*31-18:1-PE and 18:1-LPE-*d*7 (Avanti Polar Lipid) were added to each sample.

### Hemolytic activity assay

Hemolytic activities of PLA_1_ (10 mg/ml), PLC (10 mg/ml), and PLD (25,000 U/ml) were analyzed on Columbia Agar with 5% sheep blood (BD). Four-microliters of the enzymes were spotted on the plates and the plates were incubated at 37°C for 24 h.

### Fluorescence microscopy

CLSM was performed as previously reported (*11, 12, 23*). Briefly, biofilms of the *S. aureus* strain MR4 were formed in BHIG supplemented with or without PLA_1_ (200 μg/ml) at 37°C for 24 h in glass-bottomed dishes. Biofilms formed on the surface of the dishes were washed once with PBS and fixed with 1% glutaraldehyde and 4% paraformaldehyde in PBS for 10 min at 25°C. After washing three-times with PBS, biofilms were stained with FM1-43, optically cleared using iCBiofilm-H1 (Tokyo Chemical Industry, Tokyo, Japan) and visualized using LSM880 with the Airyscan high resolution module (Zeiss, Oberkochen, Germany) or THUNDER Imaging System (Leica Microsystems, Wetzlar, Germany).

### TEM

TEM analysis of *S. aureus* strain MR23 biofilms was performed as previously reported (*21*). Biofilms were grown in BHIG at 37°C for 4 h in 35-mm plastic dishes (Nunc, Roskilde, Denmark), then scraped and collected by centrifugation. The biofilm cells were fixed with 2.5% glutaraldehyde in 0.1 M phosphate buffer (pH 7.4) at 25°C for 1 h, followed by fixation with 1% osmium tetroxide in 0.1 M phosphate buffer (pH 7.4) at 4°C for 1 h. Samples were then dehydrated in an ethanol gradient at 25°C, embedded in Epon812, and thin-sectioned using an Ultracut UCT ultramicrotome (Leica, Wetzlar, Germany). Sections were stained with uranyl acetate and lead citrate and observed using an H7600 TEM (Hitachi, Tokyo, Japan) at 80 kV.

### ASEM

We used ASEM (JASM-6200, JEOL, Tokyo, Japan) to directly visualize extracellular structures within biofilms under aqueous conditions (*11, 21, 25*). *S. aureus* MR23 biofilms were grown in BHIG at 37°C for 4 h in ASEM dishes (35 mm in diameter, JEOL, Tokyo, Japan). After fixing with 1% glutaraldehyde in double distilled water (hereafter DDW) at 25°C for 10 min, the reaction was quenched by washing the specimens in 50 mM ammonium chloride followed by an additional washing in DDW. Subsequently, biofilms were labelled with positively charged nanogold (PCG) (Nanoprobes). The specimens were then washed in DDW, and the size of the gold nanoparticles was increased by gold enhancement (GoldEnhance, Nanoprobes) for 10 min at 25°C, followed by an additional washing in DDW. The resultant specimens were then incubated for 2 h at 25°C in 2 (v/v) phosphotungstic acid (TAAB Laboratories Equipment, Aldermaston, Berks, England) in DDW. After a final DDW-wash, specimens were soaked in 1% (v/v) ascorbic acid in DDW and observed using ASEM at an acceleration voltage of 30 kV.

Similarly, aggregates of strain MR4 were analyzed using ASEM.

### FE-SEM

We utilized FE-SEM to observe extracellular structures in *S. aureus* MR4 biofilms and aggregates at high resolution. Biofilms of MR4 were cultivated in BHIG medium with or without 200 μg/ml PLA_1_ at 37°C for 24 h. The biofilms were scraped into test tubes and pre-fixed overnight at 4°C in a solution containing 1.2% glutaraldehyde and 0.1 M phosphate buffer (pH 7.0). After washing with 0.1 M phosphate buffer (pH 7.0), biofilm cells were placed on 0.1% poly-lysine-coated glass pieces (5 mm × 5 mm). The cells were incubated for 30 min at 25°C in a humid chamber and then washed with 5% sucrose in 0.1 M phosphate buffer (pH 7.0) at 4°C. The cells were post-fixed with 1% OsO4 in 0.1 M phosphate buffer (pH 7.0) at 4°C for 1 h. Next, the biofilm cells were sequentially dehydrated in 50%, 70%, and 80% ethanol at 4°C for 10 min each, followed by 80%, 90%, and 100% ethanol at 25°C for 10 min. After replacing the 100% ethanol with 3-methylbutyl acetate and incubating it at 25°C for 5 min, critical point drying was performed. The specimens were then mounted on sample stages, coated with a ∼10 nm layer of osmium, and imaged using an FE-SEM (Regulus 8100, Hitachi, Tokyo, Japan). Aggregates of *S. aureus* formed in the presence of Lys-PG (C_18:1_) were prepared following the same fixation, dehydration, and osmium tetroxide (OsO_4_) coating protocols and subsequently analyzed using FE-SEM.

### Cell aggregation assay

*S. aureus* MR4 cells were cultured overnight in BHI at 37°C. The overnight culture (4 ml) was pelleted by centrifugation at 5,000 × g for 10 min at 4°C, washed three times with 1 ml of PBS, and resuspended in PBS. A 100 μl aliquot of the bacterial suspension was then mixed with the indicated concentrations of phospholipids in 1.5 ml plastic tubes. For light microscopy, a 10 μl aliquot of the bacterial suspension was placed on a glass slide and mixed with the specified concentrations of phospholipids, 1 M NaCl, and/or 200 μg/ml PLA_1_ to observe cell aggregation. For ASEM analysis, a 100 μl aliquot of the bacterial suspension was placed on an ASEM dish and incubated for at least 30 min at 25°C. Bacterial cells and aggregates were then fixed, stained, and imaged using ASEM as described above.

Additionally, bacterial cell aggregation was evaluated by light scattering. MR4 cells washed with PBS were incubated at 25°C in PBS, with or without 50 μg/ml Lys-PG (C_18:1_). The light scattering of MR4 cells at 600 nm was measured using a fluorescent spectrophotometer F7000 (Hitachi).

### Silica-Coated Microbeads Aggregation Assay

A 1 ml suspension of silica-coated microbeads (Silica Microspheres with a diameter of 0.90 μm, Polysciences, Warrington, PA, USA) was washed twice with PBS and resuspended in 1 ml of PBS. The beads were incubated with or without 100 μg/ml Lys-PG (C_18:1_) for 30 min at 25°C. When necessary, 1 M NaCl and 200 μg/ml PLA_1_ were added to the suspension. After incubation, bead aggregation was observed under a microscope using a 10× objective lens.

### Statistical analysis

Statistical analyses were performed using GraphPad Prism (version 9, GraphPad Software, Inc., San Diego, CA, USA) on Windows OS. Statistical significance was determined using one-way ANOVA with Dunnett’s or Sidak’s corrections, or two-way ANOVA with Dunnett’s correction, depending on the context. All experiments were conducted independently at least twice to ensure the accuracy and reproducibility of the findings. A *p*-value of less than 0.05 was considered statistically significant for all analyses.

## ACKNOWLEDGEMENTS

The authors would like to acknowledge Mrs. Mari Sato, Reina Miyakawa, and Naoko Toda for their experimental support. We also thank Dr. Kenji Kurokawa for providing the RN4220 mutant strains and plasmids, Dr. Taeok Bae for supplying pKOR1, and Drs. Jean-Marc Ghigo, Christophe Beloin, and Yutaka Yoshii for their critical review of the manuscript. This work was partially supported by a Grant-in-Aid for Young Scientists (A) from JSPS (no. JP15H05619 to S.S.), a Grant-in-Aid for Scientific Research (B) from JSPS (nos. JP20H02904 and JP24K01671 to S.S.), JST ERATO (no. JPMJER1502 to S.S.), and AMED-CREST (no. JP23gm1210013 to M.M.).

The authors declare no conflict of interests.

## Author contributions

S.S. planned the project. S.S., Y.T., Y.N. designed the experiments. S.S., K.H., Y.T., Y.N., and C.S. performed the experiments and analyzed the data. C.S. developed Nanogold- and heavy metal-labeling for biofilms and assisted during ASEM analysis. S.S. wrote the paper with input from all coauthors.

## References

1. H. C. Flemming, J. Wingender, The biofilm matrix. Nat Rev Microbiol 8, 623–633 (2010).

2. V. Gueneau et al., Positive biofilms to guide surface microbial ecology in livestock buildings. Biofilm 4, 100075 (2022).

3. D. Xu, T. Gu, D. R. Lovley, Microbially mediated metal corrosion. Nat Rev Microbiol 21, 705–718 (2023).

4. O. Ciofu, C. Moser, P. O. Jensen, N. Hoiby, Tolerance and resistance of microbial biofilms. Nat Rev Microbiol 20, 621–635 (2022).

5. K. Schilcher, A. R. Horswill, Staphylococcal Biofilm Development: Structure, Regulation, and Treatment Strategies. Microbiol Mol Biol Rev 84, (2020).

6. C. von Eiff, K. Becker, K. Machka, H. Stammer, G. Peters, Nasal carriage as a source of Staphylococcus aureus bacteremia. Study Group. N Engl J Med 344, 11–16 (2001).

7. W. van Schaik, T. Abee, The role of sigmaB in the stress response of Gram-positive bacteria -- targets for food preservation and safety. Curr Opin Biotechnol 16, 218–224 (2005).

8. S. Sugimoto et al., Staphylococcus epidermidis Esp degrades specific proteins associated with Staphylococcus aureus biofilm formation and host-pathogen interaction. J Bacteriol 195, 1645–1655 (2013).

9. A. Chiba, S. Sugimoto, F. Sato, S. Hori, Y. Mizunoe, A refined technique for extraction of extracellular matrices from bacterial biofilms and its applicability. Microb Biotechnol 8, 392–403 (2015).

10. S. Sugimoto et al., Broad impact of extracellular DNA on biofilm formation by clinically isolated Methicillin-resistant and -sensitive strains of Staphylococcus aureus. Sci Rep 8, 2254 (2018).

11. K. Yonemoto et al., Redundant and Distinct Roles of Secreted Protein Eap and Cell Wall-Anchored Protein SasG in Biofilm Formation and Pathogenicity of Staphylococcus aureus. Infect Immun 87, (2019).

12. A. Chiba et al., Staphylococcus aureus utilizes environmental RNA as a building material in specific polysaccharide-dependent biofilms. NPJ Biofilms Microbiomes 8, 17 (2022).

13. J. Kitaura, M. Murakami, Positive and negative roles of lipids in mast cells and allergic responses. Curr Opin Immunol 72, 186–195 (2021).

14. A. Sahebkar, Fat lowers fat: purified phospholipids as emerging therapies for dyslipidemia. Biochim Biophys Acta 1831, 887–893 (2013).

15. R. P. Rehal et al., The influence of mild acidity on lysyl-phosphatidylglycerol biosynthesis and lipid membrane physico-chemical properties in methicillin-resistant Staphylococcus aureus. Chem Phys Lipids 206, 60–70 (2017).

16. C. Wolk et al., Phase Diagram for a Lysyl-Phosphatidylglycerol Analogue in Biomimetic Mixed Monolayers with Phosphatidylglycerol: Insights into the Tunable Properties of Bacterial Membranes. Chemphyschem 21, 702–706 (2020).

17. G. P. Dubey, S. Ben-Yehuda, Intercellular nanotubes mediate bacterial communication. Cell 144, 590–600 (2011).

18. S. N. Wai et al., Vesicle-mediated export and assembly of pore-forming oligomers of the enterobacterial ClyA cytotoxin. Cell 115, 25–35 (2003).

19. Y. Tashiro, H. Uchiyama, N. Nomura, Multifunctional membrane vesicles in Pseudomonas aeruginosa. Environ Microbiol 14, 1349–1362 (2012).

20. H. C. Flemming et al., The biofilm matrix: multitasking in a shared space. Nat Rev Microbiol 21, 70–86 (2023).

21. S. Sugimoto et al., Imaging of bacterial multicellular behaviour in biofilms in liquid by atmospheric scanning electron microscopy. Sci Rep 6, 25889 (2016).

22. T. Kunoh et al., Polyfunctional Nanofibril Appendages Mediate Attachment, Filamentation, and Filament Adaptability in Leptothrix cholodnii. ACS Nano 14, 5288–5297 (2020).

23. S. Sugimoto, Y. Kinjo, Instantaneous Clearing of Biofilm (iCBiofilm): an optical approach to revisit bacterial and fungal biofilm imaging. Commun Biol 6, 38 (2023).

24. S. Sugimoto et al., Cloning, expression and purification of extracellular serine protease Esp, a biofilm-degrading enzyme, from Staphylococcus epidermidis. J Appl Microbiol 111, 1406–1415 (2011).

25. H. Nishiyama et al., Reprint of: Atmospheric scanning electron microscope observes cells and tissues in open medium through silicon nitride film. J Struct Biol 172, 191–202 (2010).

26. J. B. Park et al., Phospholipase signalling networks in cancer. Nat Rev Cancer 12, 782–792 (2012).

27. N. Liu et al., Immobilization of Lecitase(R) Ultra onto a novel polystyrene DA-201 resin: characterization and biochemical properties. Appl Biochem Biotechnol 168, 1108–1120 (2012).

28. R. M. Corrigan, D. Rigby, P. Handley, T. J. Foster, The role of Staphylococcus aureus surface protein SasG in adherence and biofilm formation. Microbiology (Reading) 153, 2435–2446 (2007).

29. M. Johnson, A. Cockayne, J. A. Morrissey, Iron-regulated biofilm formation in Staphylococcus aureus Newman requires ica and the secreted protein Emp. Infect Immun 76, 1756–1765 (2008).

30. A. Peschel et al., Staphylococcus aureus resistance to human defensins and evasion of neutrophil killing via the novel virulence factor MprF is based on modification of membrane lipids with l-lysine. J Exp Med 193, 1067–1076 (2001).

31. N. Ichihashi, K. Kurokawa, M. Matsuo, C. Kaito, K. Sekimizu, Inhibitory effects of basic or neutral phospholipid on acidic phospholipid-mediated dissociation of adenine nucleotide bound to DnaA protein, the initiator of chromosomal DNA replication. J Biol Chem 278, 28778–28786 (2003).

32. R. M. Golla et al., Resistome of Staphylococcus aureus in Response to Human Cathelicidin LL-37 and Its Engineered Antimicrobial Peptides. ACS Infect Dis 6, 1866–1881 (2020).

33. A. K. Foreman-Wykert, J. Weiss, P. Elsbach, Phospholipid synthesis by Staphylococcus aureus during (Sub)Lethal attack by mammalian 14-kilodalton group IIA phospholipase A2. Infect Immun 68, 1259–1264 (2000).

34. T. Bae, O. Schneewind, Allelic replacement in Staphylococcus aureus with inducible counter-selection. Plasmid 55, 58–63 (2006).

35. R. P. Novick et al., Synthesis of staphylococcal virulence factors is controlled by a regulatory RNA molecule. EMBO J 12, 3967–3975 (1993).

36. C. L. Laut et al., Bacillus anthracis Responds to Targocil-Induced Envelope Damage through EdsRS Activation of Cardiolipin Synthesis. mBio 11, (2020).

37. Y. Taketomi et al., Lipid-orchestrated paracrine circuit coordinates mast cell maturation and anaphylaxis through functional interaction with fibroblasts. Immunity 57, 1828–1847 e1811 (2024).

38. K. Yamamoto et al., Secreted Phospholipase A(2) Specificity on Natural Membrane Phospholipids. Methods Enzymol 583, 101–117 (2017).

