## Supplementary Materials for "Lysyl-phosphatidylglycerol promotes cell-to-cell interaction and biofilm formation of *Staphylococcus aureus* as a biofilm matrix component"

**This PDF file includes:**

Figure S1 to S6
Table S1 to S2
SI References

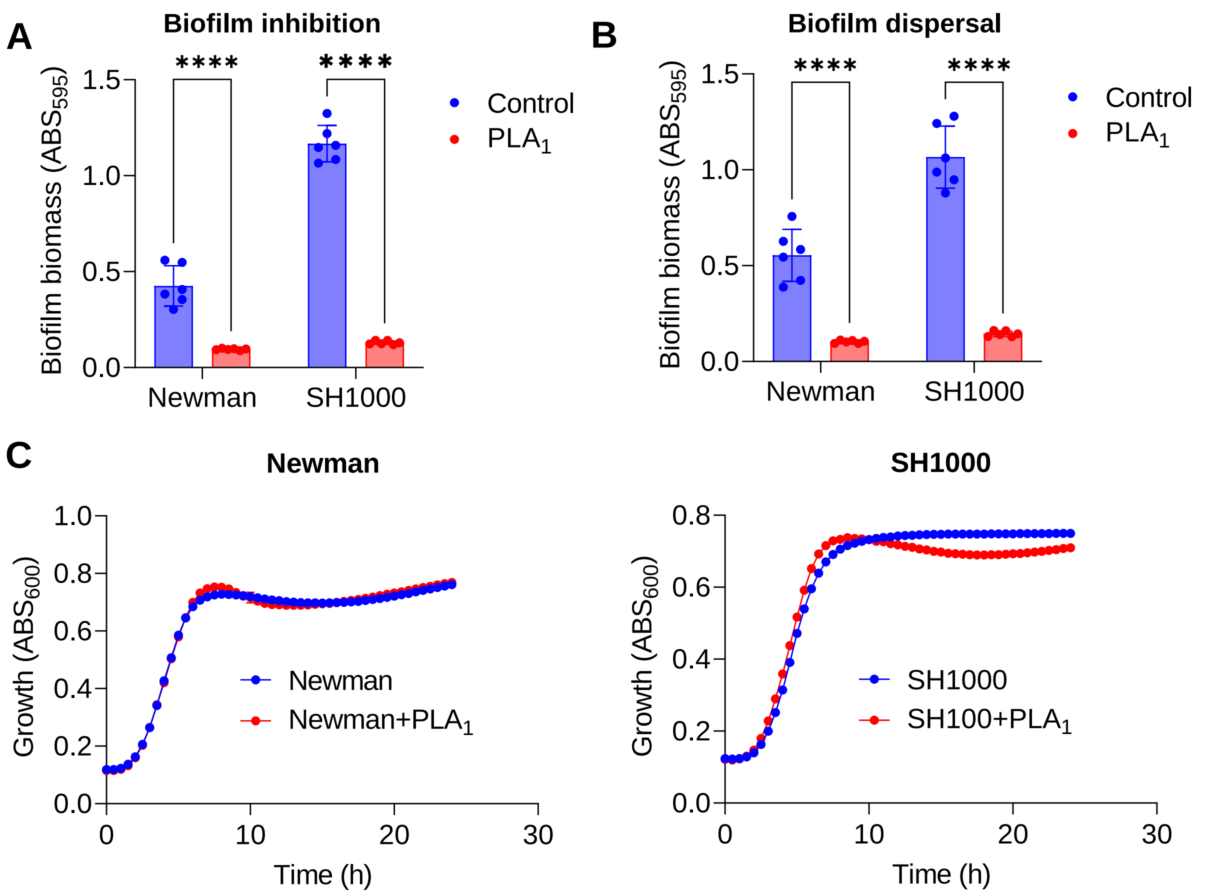


**Fig. S1. Anti-biofilm Activity of PLA1 on *S. aureus* Laboratory Strains.**

(**A**) Biofilms of the widely studied *S. aureus* strains SH1000 and Newman were grown in brain heart infusion supplemented with 1% glucose (BHIG) at 37°C for 24 h, either without treatment (Control) or in the presence of 200 µg/ml PLA$_1$. Biofilm biomass was quantified as described in Fig. 2. (**B**) The dispersal effects of 200 µg/ml PLA$_1$ on preformed biofilms were evaluated. (**C**) The growth of SH1000 and Newman strains in BHIG, with and without 200 µg/ml PLA$_1$, was monitored by measuring culture absorbance at 600 nm every 30 min over a 24-h period. Data represent means ± standard deviations from six independent experiments. Statistical significance was assessed using two-way ANOVA with Sidak's correction for multiple comparisons relative to the Control in panels (**A**) and (**B**). ****, $p < 0.0001$.

**PG ($C_{18:1}$)**

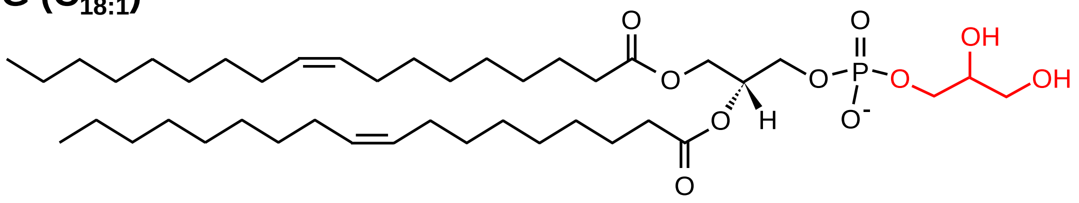

**Lys-PG ($C_{18:1}$)**

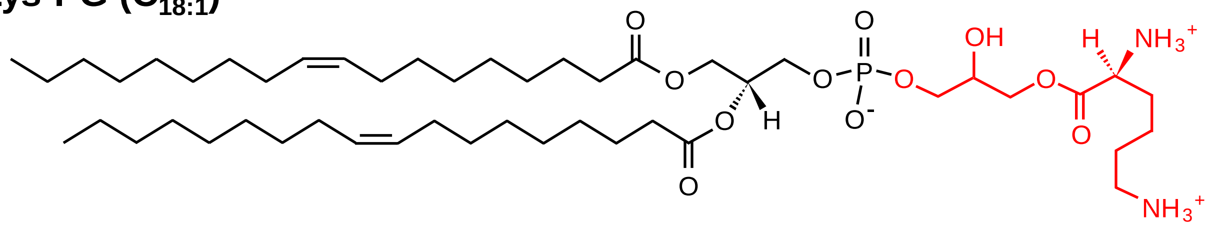

**CL ($C_{18:1}$)**

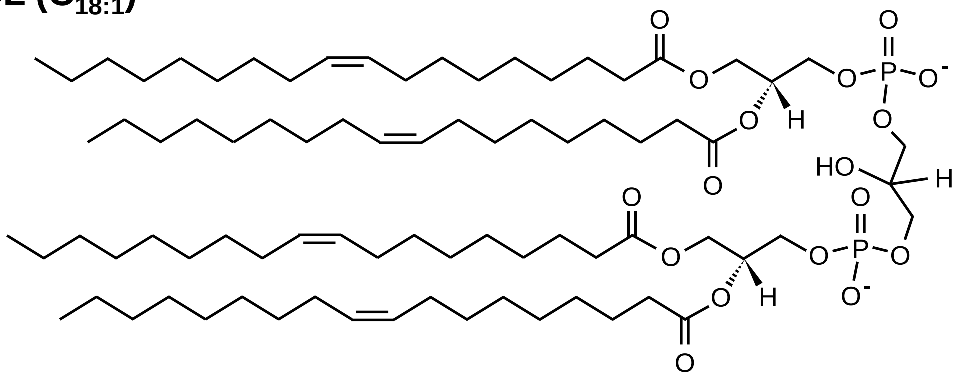

**Fig. S2. Chemical structures of the major phospholipids in *S. aureus*.** Polar head groups are colored in red.

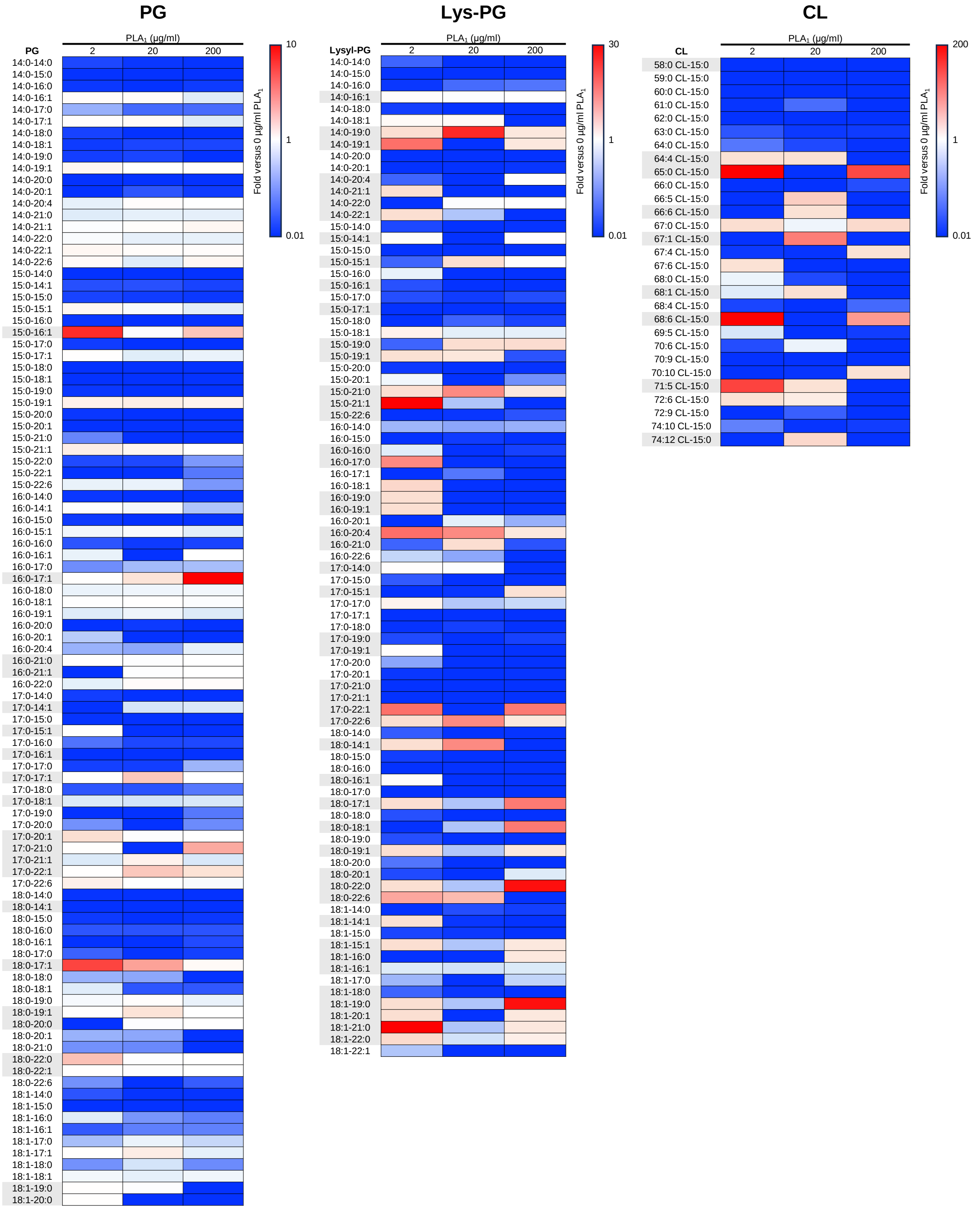


**Fig. S3. Phospholipid Cleavage Assessed by Lipidomic Analysis.**

Relative levels of major phospholipid species in the biofilm matrix of *S. aureus* MR4 were measured after treatment with varying concentrations of $PLA_1$. Data are presented as ratios to pre-treatment levels, with the abundance of each phospholipid species prior to $PLA_1$ treatment set to 1. Phospholipid species with low MS signal intensities ($<10^6$ for PG and CL; $<10^5$ for Lys-PG) were excluded from the analysis shown in Fig. 4, G to H. I.

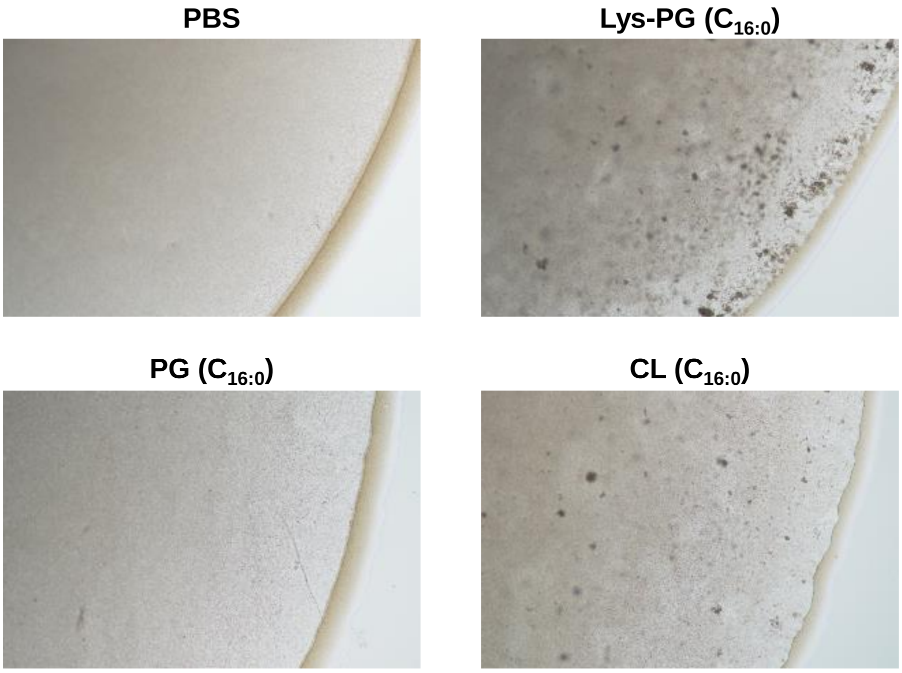


**Fig. S4. Impact of Phospholipids ($C_{16:0}$) on *S. aureus* Cell Aggregation.**

*S. aureus* MR4 cells were incubated with 100 µg/ml of PG ($C_{16:0}$), Lys-PG ($C_{16:0}$), or CL ($C_{16:0}$) in phosphate-buffered saline (PBS). Following a 30-min incubation at 25°C, 10 µl aliquots were placed on a slide glass for direct observation of cell aggregation using optical microscopy with a 10× objective lens. A control sample without phospholipids in PBS was included for comparison.

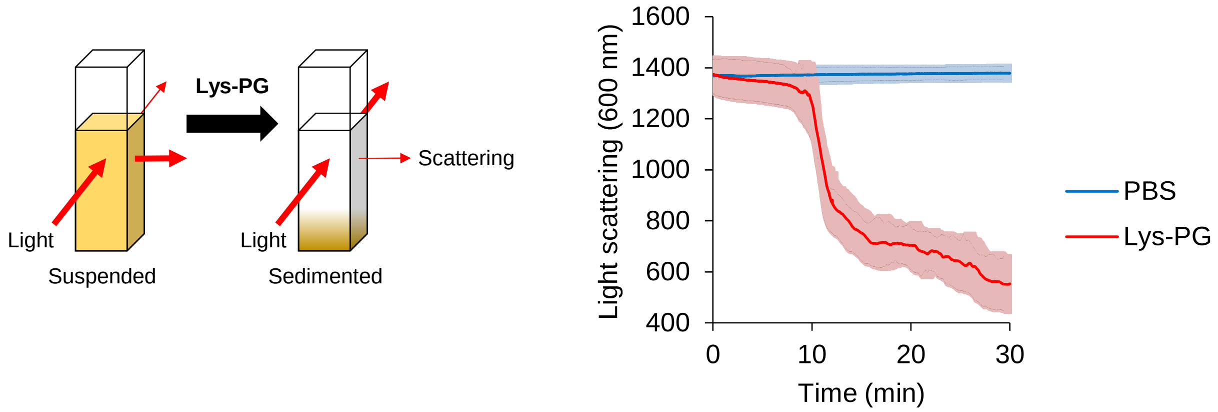


**Fig. S5. Time-Dependent Aggregation of *S. aureus* Induced by Lys-PG.**

The aggregation of *S. aureus* MR4 cells, resulting in sedimentation, was assessed by monitoring light scattering over time. Cells were incubated at 25°C in PBS in the presence or absence of 50 μg/ml Lys-PG ($C_{18:1}$). Data represent the means and standard deviations from three independent measurements.

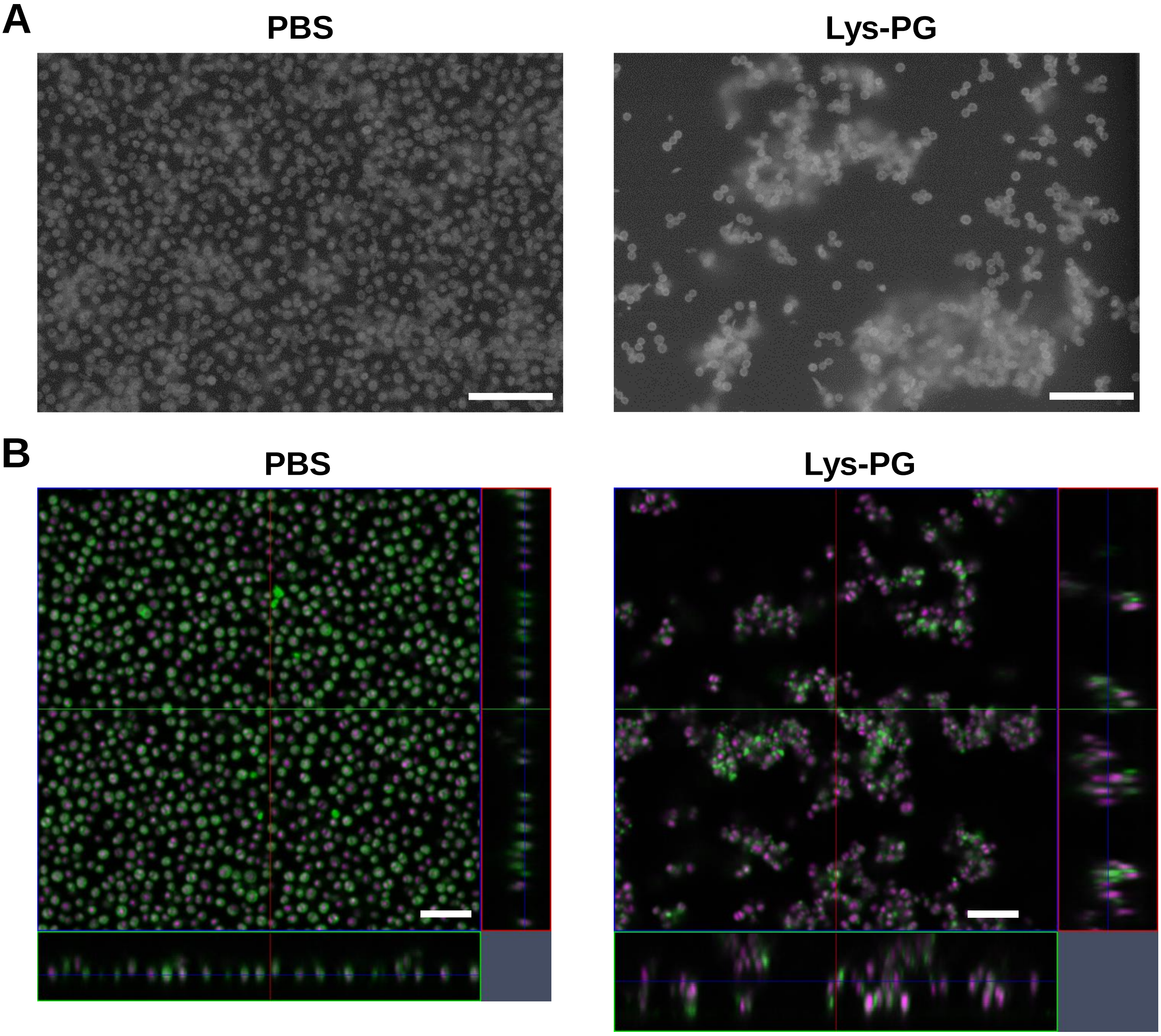


**Fig. S6. Visualization of *S. aureus* Cell Aggregation and Surface Attachment.**

(**A**) Atmospheric scanning electron microscopy (ASEM) imaging of cell aggregation and attachment. *S. aureus* MR4 cells were incubated with or without 100 µg/ml Lys-PG ($C_{18:1}$) in PBS on ASEM dishes. Following 30 min of incubation at 25°C, cells were fixed with 1% glutaraldehyde, stained with positively charged nanogold and phosphotungstic acid, and visualized using ASEM. Representative images are shown. Scale bar: 10 µm. (**B**) CLSM imaging of cell aggregation and attachment. *S. aureus* MR4 cells were incubated in PBS containing 100 µg/ml Lys-PG ($C_{18:1}$) on glass-bottom dishes. Control samples lacked Lys-PG ($C_{18:1}$) supplementation. After 30 min of incubation at 25°C, cells were fixed with 1% glutaraldehyde, stained with FM1-43 (membrane marker) and DRAQ5 (DNA marker), and imaged using an LSM880 microscope equipped with an Airyscan super-resolution unit. Representative images are presented. Scale bar: 5 µm.

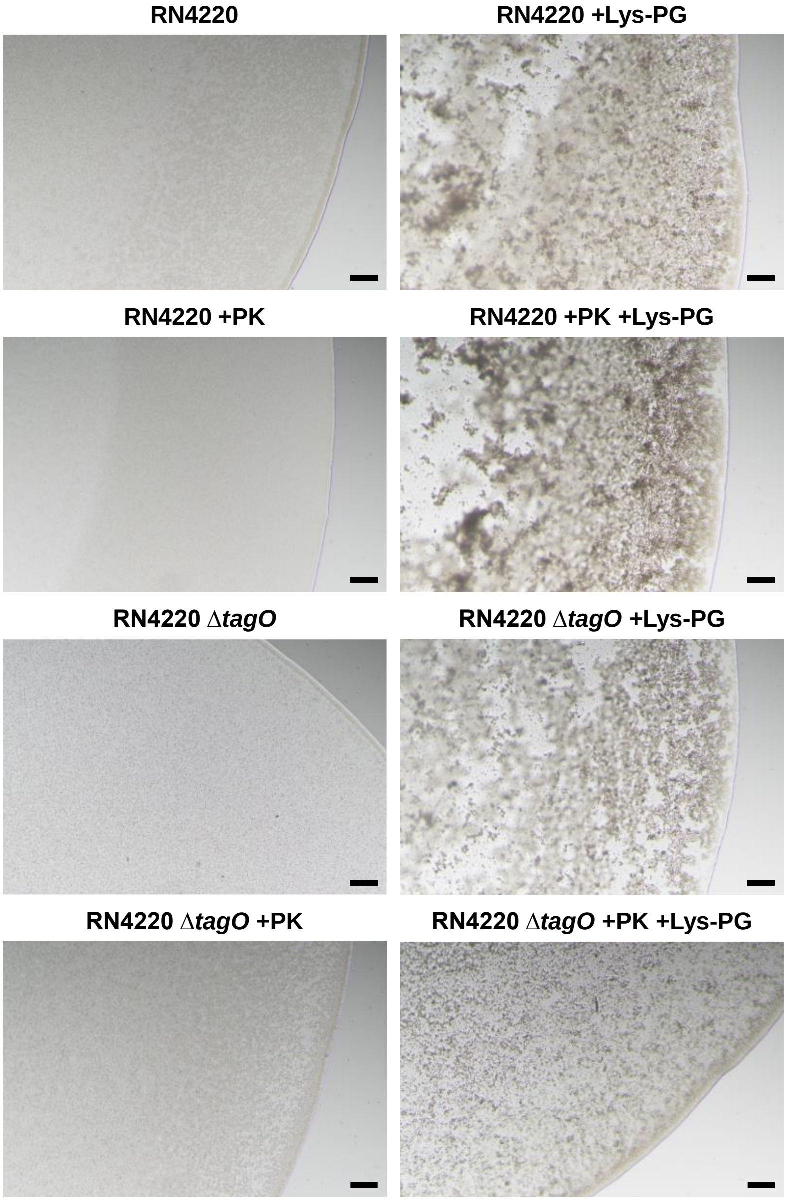


**Fig. S7. Influence of Cell Surface Molecules on Lys-PG-Induced Aggregation of *S. aureus*.**
*S. aureus* RN4220 and its derivative strains were grown overnight in 5 ml of BHI at 37°C. The overnight cultures (4 ml) were washed three times with PBS and resuspended in PBS (4 ml). Where indicated, cells were treated with 200 µg/ml proteinase K (PK) at 37°C for 2 h to digest cell surface proteins before being washed again with PBS. The resulting suspensions were then mixed with or without 100 µg/ml Lys-PG ($C_{18:1}$) in test tubes. Small aliquots (10 µl) of the mixtures were placed on slide glasses and incubated at 25°C for 30 min. Following incubation, cell aggregates were visualized using optical microscopy with a 10× objective lens. Scale bar: 100 µm.

**Table S1. Bacterial strains and plasmids used in this study.**

| Strains/plasmids | Description | Source/Reference |
|---|---|---|
| *S. aureus* | | |
| RN4220 | NCTC8325-4 derivative, restriction deficient mutant | (*1*) |
| SH1000 | NCTC8325-4 with functional *rsbU* | (*2*) |
| M0702 | RN4220 *tagO*::pT0702 | (*3*) |
| Newman | Laboratory strain, high level of clumping factor | (*4*) |
| MR4 | A clinical isolate of MRSA from the Jikei hospital | (*5*) |
| MR4 Δ*mprF* | *mprF* was deleted from MR4 | This study |
| MR11 | A clinical isolate of MRSA from the Jikei hospital | (*5*) |
| MR23 | A clinical isolate of MRSA from the Jikei hospital | (*6*) |
| *E. coli* | | |
| DH5α | *fhuA2* Δ(*argF*-*lacZ*)*U169 phoA glnV44 φ80* Δ(*lacZ*)*M15 gyrA96 recA1 relA1 endA1 thi-1 hsdR17* | Toyobo |
| Plasmids | | |
| pHY300PLK | A shuttle vector between *E. coli* and *S. aureus*, $Tet^R$ | Takara |
| pHYmprF | *mprF* from RN4220 was cloned into pHY300PLK | (*7*) |
| pKOR1 | *E. coli*-*S. aureus* shuttle vector plasmid for knockout of genes by allelic exchange, $Amp^R$, $Cm^R$ | (*8*) |
| pMprF-KO | pKOR1-derivative plasmid for knockout of *mprF* in MR4, $Amp^R$, $Cm^R$ | This study |

$Amp^R$, ampicillin-resistant; $Cm^R$, chloramphenicol-resistant; $Tet^R$, tetracycline-resistant.

**Table S2. Oligonucleotide primers used in the current study.**

| Primers | Sequences (5' to 3') | Descriptions |
|---|---|---|
| attB1-mprF-F | GGGGACAAGTTTGTACAAAAAAGCAGGCTGTAATAAAATAGTTGAATAAGTAATAAAAAATACCAATGAC | Disruption of *mprF* in MR4 |
| mprF-R | GCACTTGGATTTTAATTTTTCACATCAATTCTAATTATTTCTGTTATAAATC | Disruption of *mprF* in MR4 |
| mprF-F | AATTGATGTGAAAAATTAAAATCCAAGTGCTAAGAGGTATACAG | Disruption of *mprF* in MR4 |
| attB2-mprF-R | GGGGACCACTTTGTACAAGAAAGCTGGGTATTAGAAATATTTTTCTCAGTCATTGACCC | Disruption of *mprF* in MR4 |

**SI References**


1. R. P. Novick *et al.*, Synthesis of staphylococcal virulence factors is controlled by a regulatory RNA molecule. *EMBO J* **12**, 3967-3975 (1993).
2. M. J. Horsburgh *et al.*, sigmaB modulates virulence determinant expression and stress resistance: characterization of a functional rsbU strain derived from *Staphylococcus aureus* 8325-4. *J Bacteriol* **184**, 5457-5467 (2002).
3. C. Kaito, K. Sekimizu, Colony spreading in *Staphylococcus aureus*. *J Bacteriol* **189**, 2553-2557 (2007).
4. E. S. Duthie, Variation in the antigenic composition of staphylococcal coagulase. *J Gen Microbiol* **7**, 320-326 (1952).
5. S. Sugimoto *et al.*, Broad impact of extracellular DNA on biofilm formation by clinically isolated Methicillin-resistant and -sensitive strains of *Staphylococcus aureus*. *Sci Rep* **8**, 2254 (2018).
6. S. Sugimoto *et al.*, Cloning, expression and purification of extracellular serine protease Esp, a biofilm-degrading enzyme, from *Staphylococcus epidermidis*. *J Appl Microbiol* **111**, 1406-1415 (2011).
7. N. Ichihashi, K. Kurokawa, M. Matsuo, C. Kaito, K. Sekimizu, Inhibitory effects of basic or neutral phospholipid on acidic phospholipid-mediated dissociation of adenine nucleotide bound to DnaA protein, the initiator of chromosomal DNA replication. *J Biol Chem* **278**, 28778-28786 (2003).
8. T. Bae, O. Schneewind, Allelic replacement in *Staphylococcus aureus* with inducible counter-selection. *Plasmid* **55**, 58-63 (2006).
